# Senescent peripheral fibroblasts initiate chemotherapy-induced peripheral neuropathy

**DOI:** 10.64898/2026.08.04.742055

**Authors:** Taylor Malachowski, Ganesh Kumar Raut, Qihao Ren, Radhika Mishra, Satarupa Mullick Bagchi, Daden Deaver, Xianmin Luo, Nathan P. Staff, Theodore J. Price, Simon Haroutounian, Sheila A. Stewart

## Abstract

Chemotherapy-induced peripheral neuropathy (CIPN) is a common, debilitating complication of cancer therapy. Although axonal degeneration defines CIPN, the initiating cellular events remain unknown. Here, we show that CIPN is initiated by senescent peripheral fibroblasts rather than by the neuron itself. Across the mechanistically distinct chemotherapeutic agents paclitaxel and cisplatin, chemotherapy-induced senescence is unexpectedly restricted to peripheral fibroblasts rather than neurons and acts upstream of neuronal SARM1 activation. Genetic or pharmacological ablation of senescent fibroblasts prevents neuropathy and reverses established disease, demonstrating that these cells are required for both disease initiation and maintenance. Mechanistically, senescent fibroblasts drive neuropathic injury through an MK2-dependent senescence-associated secretory phenotype (SASP), and genetic or pharmacological inhibition of MK2 suppresses the SASP, preserves peripheral innervation and restores sensory function. Together, these findings redefine the cellular origin of CIPN and identify MK2-dependent fibroblast senescence as a therapeutic target.

## INTRODUCTION

Chemotherapy is a mainstay of cancer therapy. In 2021, approximately 1.9 million new cancer cases were diagnosed and ∼57% required chemotherapy treatment [1]. While chemotherapy can profoundly impact disease free survival, it is unfortunately often accompanied with devastating side effects including chemotherapy induced peripheral neuropathies (CIPN) [2]. CIPN is a particularly insidious side effect as it can persist in over 30% of patients. CIPN is observed in patients receiving cytotoxic agents such as platinum compounds, taxanes, and vinca alkaloids. Permanent symptoms can include tactile or thermal allodynia (pain), numbness, and dysesthesia (tingling) in the hands and feet [3]. Currently, there are no preventative or curative treatments for CIPN [4], and patients are relegated to treatments that only dampen symptoms [3]. Thus, the only approach to mitigating CIPN is dose reduction or cessation of chemotherapy, which can significantly impact patient survival [1]. Given the prevalence and lack of treatment options, a detailed understanding of the biological mechanisms underlying CIPN is of the upmost importance.

Axons ensure proper nerve function by creating long-distance connections between neurons and distal tissues. Axon terminals that innervate distal tissues form specialized structures called nociceptors or mechanoreceptors, playing a central role in sensory perception and responses to stimuli. Structural and functional defects in these connections, known as axonopathies, can severely compromise function [5]. Toxins, direct injuries, and diseases trigger signaling cascades that can lead to axon destruction and nerve dysfunction. Taxanes, commonly used to treat cancers like breast cancer, can directly affect neurons by disturbing mitochondrial function, calcium signaling, and axonal transport [6]. Axon degeneration, induced by taxanes, starts at the terminals and progresses retrogradely towards the neuron cell body [7] and is commonly observed in CIPN [6]. While it remains an open question whether toxins and mechanical injury use similar mechanisms to initiate axonal degradation, emerging data suggest they converge on common pathways within the neuron. Recent work from DiAntonio, Millbrandt, and colleagues demonstrated that injury and CIPN activate neuronal SARM1 that drives axonal degradation [8]. However, what drives neuronal SARM activation remains an open question.

Paclitaxel (PTX) impacts cells throughout the body by inducing several cellular changes including DNA damage and, in some settings, cellular senescence. Senescent cells are typically characterized by cell cycle arrest, increased CDKN2a (i.e., p16) expression, increased senescence associated β-galactosidase (SA-β-gal) hydrolyzation, apoptosis resistance, and expression of the senescence-associated secretory phenotype (SASP) factors [9] including IL-6 and TNF-α that have been implicated in nerve homeostasis [10–12]. Here we find that PTX drives p38MAPK-MK2 dependent SASP in peripheral PDGFR ⍰^+^ fibroblasts that drives axonal denervation and CIPN. Further, we show that senolytics or an MK2 inhibitor can prevent and even reverse PTX driven CIPN, raising the possibility that we can treat patients suffering from CIPN and return quality of life.

## RESULTS

### PTX reduces primary tumor growth while inducing denervation and symptoms of CIPN

PTX remains a frontline therapy for many cancer types including breast, ovarian and lung cancers where it is deployed in both early stage and advanced cases. Unfortunately, PTX can also induce CIPN in 61-92% of patients depending on the dose [13]. The development of CIPN symptoms such as pain or numbness in the hands and feet can led to dose de-escalation and in some cases, cessation of treatment resulting in disease progression. To establish a model for CIPN, we injected 1×10^5^ PyMT-Bo-1 (GFP/Luc) breast cancer cells into the mammary fat pad of 12-week-old female C57BL/6J mice to establish a primary breast tumor. After confirmation at Day 7 of tumor sizes greater than 100mm^3^ via caliper measurement, mice were treated with Vehicle (Veh), or PTX (50 mg/kg, 2 doses) via tail vein injection. Mice were tested for symptoms of sensory neuropathy via the Von Frey test. Baseline Von Frey responses were collected one day before PTX treatment, and post-treatment responses were collected one day before sacrifice. On day 9 mice were sacrificed and tumor growth and axon innervation in the plantar surface of the hind paw was assessed (**Fig. 1A**). As expected, PTX significantly reduced tumor growth (**Fig. 1B**). To assess neuronal innervation, we performed immunofluorescent (IF) staining of the pan-axonal marker protein gene product 9.5 (PGP9.5) to label peripheral nervous system (PNS) axons on paw tissue and quantitated axon innervation [14]. We found that PTX treatment led to significant axon loss (**Fig. 1C-D**). In addition, when comparing baseline Von Frey measurements, to those at endpoint, mice treated with PTX exhibited an increased Paw Withdraw Threshold (PWT), indicating that animals required more force to elicit a response to stimuli and were experiencing numbness (**Fig 1E**).

**Figure 1:**
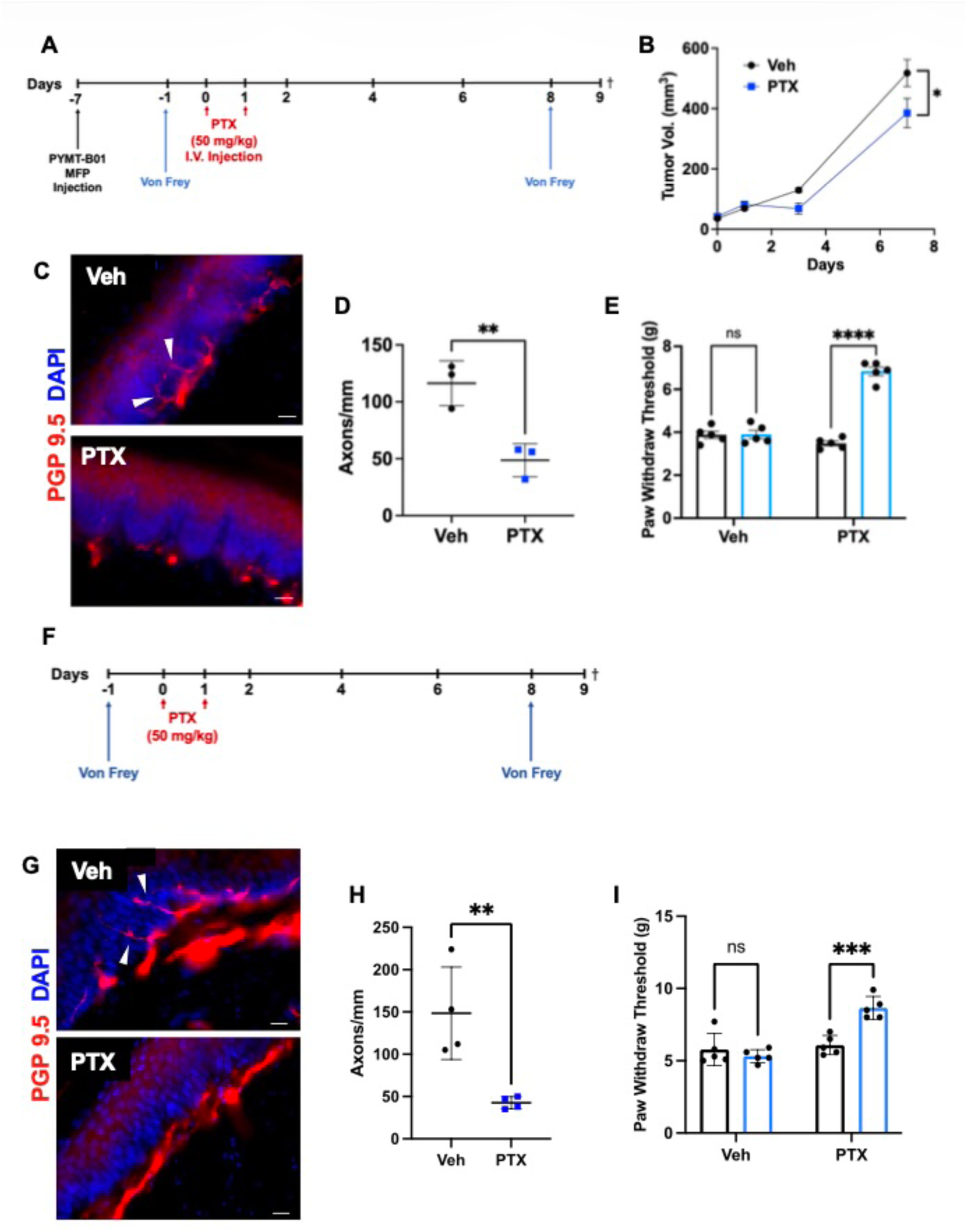
PTX reduces primary tumor growth while inducing axonal drawback and symptoms of CIPN. **(A)** Schema: PTX treatment delivery in mice with MFP injection of PYMT-Bo1 tumor cells. **(B)** Tumor growth was assessed by caliper measurement on indicated time points (2-way ANOVA was performed to compare the tumor growth; data are represented as mean ± SEM. *p<0.05). **(C)** Representative immunofluorescence (IF) of PGP9.5 (red) and DAPI (blue) in hind paw tissues from Veh or PTX treated mice. **(D)** Quantification of innervating axons from same mice shown in C; (Student t-test was performed to compare innervating axon numbers, data are represented as mean ± SEM. **p<0.01). **(E)** Von Frey results on Day -1 (black) and Day 8 (blue) from same mice shown in C; (2-way ANOVA was performed to compare behavioral outputs, data are represented as mean ± SEM. ***p <0.001; ns, not significant). **(F)** Schema: PTX treatment delivery in non-tumor bearing WT mice. **(G)** Representative IF images of PGP9.5 (red) and DAPI (blue) in hind paw tissues from Veh or PTX treated mice from experimental timeline shown in F. **(H)** Quantification of innervating axons from same mice shown in G; (Student t-test was performed to compare innervating axon numbers, data are represented as mean ± SEM. **p<0.01). **(I)** Von Frey results on Day -1 (black) and Day 8 (blue) from same mice shown in G; (2-way ANOVA was performed to compare behavioral outputs, data are represented as mean ± SEM. ***p <0.001; ns, not significant).

To establish a simplified system to study CIPN, we treated wildtype (WT) C57Bl/6 mice without tumors with PTX and 9 days later assessed axon innervation and behavioral readouts. Similar to the tumor model, a baseline Von Frey assay was conducted 1 day prior to PTX treatment and post-treatment responses were collected 1 day before sacrifice (**Fig. 1F**). Similar to tumor bearing animals, we observed significant axon withdraw (**Fig. 1G-H**) and an increase in the PWT in animals treated with PTX (**Fig. 1I**).

### Genetic and pharmacologic elimination of senescent cells prevents CIPN

Chemotherapy induces a panoply of effects beyond tumor cell death including the induction of senescence. Because senescence can have a wide range of effects on tissue homeostasis and senescence associated secretory phenotype factors (SASP) including IL6 and TNFα have been implicated in CIPN [10, 11], we asked if it contributed to CIPN. To address this important question, we utilized INK-ATTAC mice that express a p16^INK4a^ dependent inducible suicide gene that allows for the selective elimination of senescent cells [15]. For these studies, 12-week-old INK^-^ and INK^+^ mice were treated with PTX followed by vehicle (Veh) or AP20187 (AP, 10mg/kg) on days 2, 4, 6, and 8 to eliminate p16^+^ senescent cells. Mice were also subjected to the Von Frey assay before treatment to establish a baseline and at endpoint to determine changes in sensory nerve function. (**Fig. 2A**). At baseline, all mice displayed similar PWTs. However, following PTX treatment, mice displayed an increase in PWT, indicating numbness. This increase was prevented upon AP treatment (**Fig. 2D**). Morphologic analysis of the axons that were visualized by staining with the pan-neuronal antibody PGP9.5 revealed that PTX reduced axon innervation, and this was inhibited when PTX treated INK^+^ mice also received AP (**Fig. 2B-C**). Together, these data indicate that senescent cells drive CIPN. Similar results were observed in male INK-ATTAC mice (**Extended Data Fig. 1A**).

**Figure 2:**
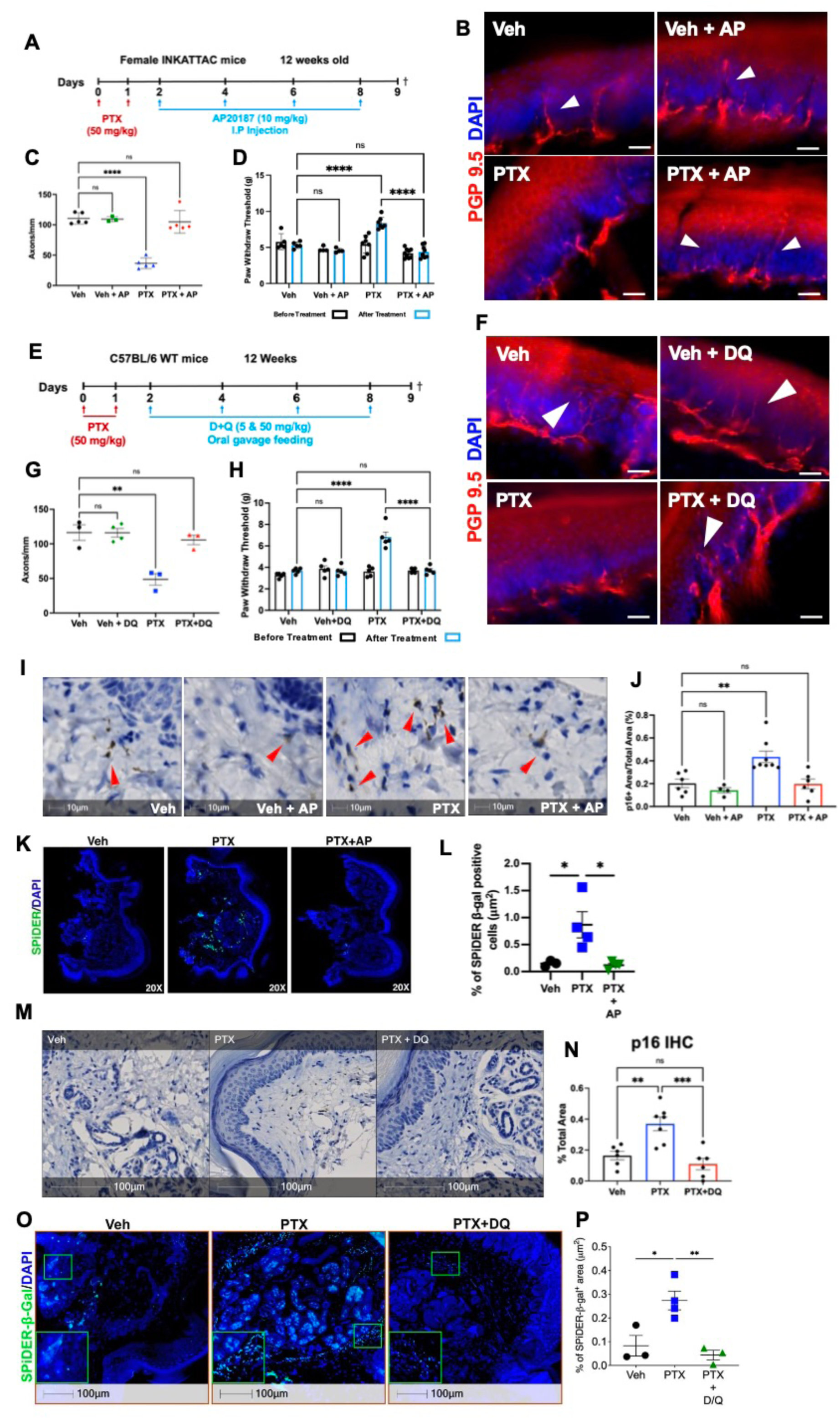
Genetic and pharmacologic elimination of senescent cells prevents CIPN. **(A)** Schema: AP treatment regimen of INK-ATTAC mice treated with Veh or PTX. **(B)** Representative IF images for PGP9.5 (red) and DAPI (blue) in hind paw sections from 12-week-old INK-ATTAC mice treated with Veh (n=5), Veh+AP (n=3), PTX (n=5) or PTX+AP (n=5). White arrowheads indicate innervating axons. **(C)** Quantification of innervating axons from same mice shown in B; (One-way ANOVA was performed, data are represented as mean ± SEM. ****p< 0.0001; ns, not significant). **(D)** Von Frey results on Day -1 (black) and Day 8 (blue) from same mice shown in C; (2-way ANOVA was performed, data are represented as mean ± SEM. ****p <0.001; ns, not significant). **(E)** Schema: Dasatinib and Quercetin (D&Q) treatment regimen for WT mice treated with Veh or PTX. **(F)** Representative IF images for PGP9.5 (red) and DAPI (blue) in hind paw sections from 12-week-old WT mice treated with Veh (n=3), Veh+DQ (n=4), PTX (n=3) or PTX+DQ (n=3). White arrowheads indicate innervating axons. **(G)** Quantification of innervating axons from same mice shown in F; (One-way ANOVA was performed, data are represented as mean ± SEM. **p< 0.01; ns, not significant). **(H)** Von Frey results on Day -1 (black) and Day 8 (blue) from same mice shown in G; (2-way ANOVA was performed, data are represented as mean ± SEM. ****p <0.001; ns, not significant). **(I)** Representative IHC images for p16 (brown) in hind paw sections from 12-week-old INK-ATTAC mice treated with Veh (n=6), Veh+AP (n=4), PTX (n=7), or PTX+AP (n=6). Red arrowheads indicate p16^+^ cells. **(J)** Quantification of p16^+^ area per total tissue area from same mice shown in I. One-way ANOVA was performed; data are represented as mean ± SEM. **p<0.01; ns, not significant). **(K)** Representative IF images for SPiDER (green) and DAPI (blue) in hind paw sections from 12-week-old INK-ATTAC mice treated with Veh (n=3), PTX (n=4) or PTX+AP (n=3). **(L)** Quantification of SPiDER-β-gal^+^ cells per total tissue area from same mice shown in K. One-way ANOVA was performed; data are represented as mean ± SEM. *p<0.05). **(M)** Representative IHC images for p16 (brown) in hind paw sections from 12-week-old WT mice treated with Veh (n=6), PTX (n=7), or PTX+DQ (n=6). **(N)** Quantification of p16^+^ area out of total tissue area from same mice shown in M. One-way ANOVA was performed, data are represented as mean ± SEM. **p< 0.01, ***p<0.001, ns, not significant. **(O)** Representative IF images for SPiDER (green) and DAPI (blue) in hind paw sections from 12-week-old WT mice treated with Veh (n=3), PTX (n=4) or PTX+DQ (n=3). **(P)** Quantification of SPiDER-β-gal^+^ cells per total tissue area from same mice shown in O. One-way ANOVA was performed; data are represented as mean ± SEM. *p<0.05, **p<0.01).

While the INK-ATTAC mouse model demonstrated that genetic elimination of p16^+^ senescent cells can prevent axon withdraw and symptoms of numbness, commonly associated with CIPN, we wanted to test the feasibility of pharmacologically targeting senescent cells to combat PTX-induced CIPN. To do so, we utilized senolytic drugs. The combination of dasatinib and quercetin (DQ) is a potent senolytic drug that kills senescent cells by poorly understood mechanisms [16]. To ask if senolytics might represent a therapeutic opportunity, 12-week-old mice were treated with Veh or PTX and on days 2, 4, 6, and 8 the mice were treated with DQ (5 & 50mg/kg) via oral gavage (**Fig. 2E**). On day 9, mice were sacrificed, and axon innervation was quantitated by PGP9.5 staining. DQ treatment alone had no impact on axon innervation in Veh treated mice, but the axon loss observed in PTX treated mice was completely inhibited in mice that also received DQ (**Fig. 2F-G**). In addition, Von Frey assays revealed that while PTX only treated animals exhibited an increased PWT, animals that also received DQ never developed symptoms of CIPN (**Fig. 2H**).

To explore if the role of senescence was restricted to PTX-induced CIPN, we went on to test another chemotherapy agent, cisplatin, in combination with DQ. Cisplatin is a platinum-based chemotherapy agent that also results in sensory-predominate neuropathy. Over 60% of patients treated with cisplatin experience symptoms of neuropathy after 1 month of treatment [17]. To establish cisplatin induced neuropathy, we treated 12-week-old female C57BL/6J mice with Veh or cisplatin (2.3mg/kg, 10 doses) for 5 days, followed by one week of rest and 5 additional days of treatment. One group of mice received DQ every other day along with cisplatin treatment. Mice received a baseline Von Frey assay one day before treatment began and a follow-up Von Frey one day before sacrifice (**Extended Data Fig. 2A**). Axon quantification revealed that cisplatin treated mice exhibited denervation that could be prevented with DQ (**Extended Data Fig. 2B**). Furthermore, mice treated with cisplatin alone displayed decreased PWTs indicative of pain, which was absent in mice treated with DQ (**Extended Data Fig. 2C**).

### Senescent cells are located in the hind paws after PTX treatment

Chemotherapy is a systemic treatment and our finding that senescence contributed to CIPN raised the important question, which cells underwent senescence in response to PTX. Given previous work suggesting cisplatin induced senescence in dorsal root ganglia (DRG) treated *in vitro*, we first assessed DRGs in our mice [18]. We found that DRGs collected from PTX treated animals displayed no changes in p16 expression (**Extended Data Fig. 3A-B**) or fluorescent SPiDER staining for SA-β-gal expression compared to Veh treated animals (**Extended Data Fig. 3C-D**). Having failed to identify senescent cells in the DRGs of PTX treated mice, we next examined cells along the length of the sural and sciatic nerves by staining for p16. We again found no changes in the number of p16^+^ cells along the sural or sciatic nerves after PTX treatment (**Extended Data Fig. 3E-H**).

Cells in the periphery can impact axonal innervation [19], so we next examined the periphery for evidence of p16 expression. First, we stained the planar surface of the hind paws and found that p16^+^ cells were elevated in PTX treated INK mice and these were lost upon AP treatment (**Fig. 2I-J**). In addition, fluorescent SPiDER staining for SA-β-gal revealed increased SPiDER^+^ cells in PTX treated hind paws (**Fig. 2K-L**). Finally, IHC for p16 and IF for SPiDER expression revealed that DQ could reduce markers of senescence after PTX treatment (**Fig. 2M-P**). Similarly, IHC p16 staining of tissue from cisplatin in treated mice revealed increased p16^+^ cells in the hind paw skin of mice treated with cisplatin that were reduced by DQ (**Extended Data Fig. 2D**). Together these findings indicated that treatment with PTX induces senescence specifically within the skin of the paws, but not within the neuronal bodies in the DRGs or the cells along the nerve.

### Senescent cells are present in human CIPN patient skin

Since we could identify senescent cells in the hind paws of mice with CIPN, we next wanted to ask if senescent cells were also present in the skin of patients suffering from CIPN. To address this, we utilized a publicly available bulk RNA-sequencing dataset from female breast cancer patients that received PTX and aged matched healthy women [20, 21]. We found that patients with CIPN displayed increased expression of Cdkn2a, the gene encoding for p16 compared to age matched controls. These patients also displayed increased expression of other senescence and SASP related genes, such as MMP13 and IL1α (**Fig 3A**). Skin punch biopsies were also available from these patients and IHC staining for p16 revealed an increase in p16^+^ cells in patients with CIPN (**Fig. 3B-C**). These results supported our mouse model findings and suggested a role for senescent cells in human CIPN.

**Figure 3:**
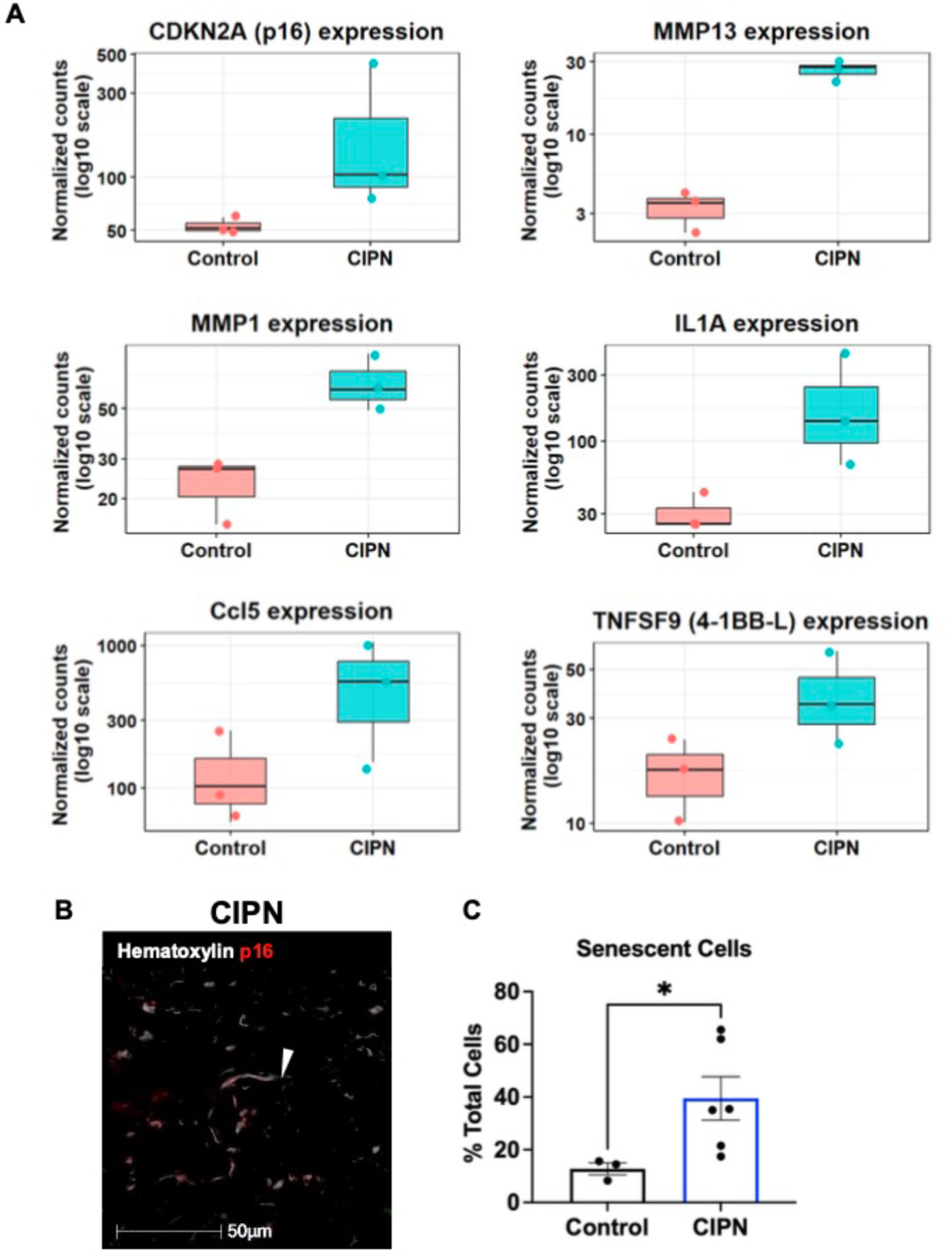
Human CIPN patients show increased senescent cells. **(A)** Box plots of gene expression between Control (n=3) or CIPN (n=3) patient samples. Genes include: *Cdkn2a*, *Mmp13*, *Mmp1*, *Il1α*, *Ccl5*, and *Tnfsf9*. **(B)** IHC were performed on skin punch biopsies from either Control (n=3) or CIPN (n=3) patients using p16 (red) and nuclei were counterstained with hematoxylin (shown as white). Cells considered p16^+^ are denoted by white arrowheads. **(C)** Quantification of senescent cells (p16^+^) out of total cells. One-tailed Student t-test was performed, data are represented as mean ± SEM. *p< 0.05.

### SARM1 activation functions downstream of senescence to drive CIPN

Recent work from DiAntonio, Milbrandt, and colleagues demonstrated that injury and CIPN activate neuronal Sterile alpha and TIR motif containing 1 (SARM1) that drives axonal degradation [8]. SARM1 is a NADase that is normally in an inactive state within neurons but is activated under conditions of stress or injury [22, 23]. In a CIPN model utilizing the chemotherapeutic agent vincristine, Geisler et al. demonstrated that loss of SARM1 prevented axonal drawback and the development of CIPN symptoms as measured by the Von Frey and cold plate assay [24]. One well studied activator of SARM is an increased ratio of nicotinamide mononucleotide (NMN) to NAD^+^ [23], but given SASP factors can impact tissue microenvironments [25], we asked if senescent fibroblast-derived SASP acted upstream of SARM1 to induce CIPN. To answer this, 12-week-old SARM1 knock out (KO) (JAX # 018069) mice were treated with PTX and assessed for axon innervation and changes in Von Frey outputs after 9 days (**Extended Data Fig 4A**). As expected, denervation and increases in PWTs in PTX treated SARM1 KO was prevented (**Extended Data Fig. 4B-C**). Despite the prevention of CIPN, mIHC analyses revealed that senescent p16^+^ cells were present in PTX treated SARM1 KO animals (**Extended Data Fig. 4D**). This suggests that the induction of senescence is upstream of SARM1 activation and subsequent axonal withdraw in PTX-induced peripheral neuropathy.

### Senescent dermal fibroblasts drive CIPN

Our p16 IHC staining and fluorescent SPiDER staining for SA-β-gal revealed that senescent cells arose at the most distal end of the nerve in the skin of the hind paws and not along the nerve or in the DRGs. Thus, we next wanted to identify the senescent cell type so that we could investigate the mechanism responsible for CIPN. Recent work by DiAntonio, Jeffrey Milbrandt, and colleagues demonstrated that macrophages are important for other forms of neuropathy [26], so we first wanted to investigate the possibility that a senescent immune cell drove CIPN in our model. To do this, we carried out a bone marrow transplant using bone marrow from INK-ATTAC mice into WT mice. This allowed us to induce the p16*^INK4a^*-driven suicide gene in senescent CD45^+^ derived cells exclusively. Recipient WT C57BL/6 (CD45.1) mice were treated with two doses of 400 cGy 4 hours apart and then received bone marrow (5×10^6^ cells/200μl/mouse) from INK-ATTAC mice (CD45.2) and animals were allowed to recover for 6 weeks (**Extended Data Fig. 5A**). To assess chimerism, peripheral blood mononuclear cells (PBMC) from recipient mice were stained with anti-CD45.2 and anti-CD45.1 antibodies and we found that mice obtained greater than 90% chimerism (**Extended Data Fig. 5B-C**). Recipient mice were then treated with PTX ± AP and 9 days later were assessed for axon innervation. We found that AP-treatment failed to prevent PTX-induced axonal withdraw, indicating that p16^+^ senescent bone marrow derived immune cells did not contribute to CIPN (**Extended Data Fig. 5D-E**).

A bone marrow transplant does not rule out the possibility of tissue resident macrophages contributing to our phenotype. To address this, we utilized a multiplex immunohistochemistry (mIHC) approach to stain INK mice treated with Veh, PTX and PTX+AP for p16^+^ F4/80^+^ macrophages. Quantification of senescent macrophages (p16+F4/80+) revealed no significant changes between groups (**Extended Data Fig. 5F-G**). These results confirmed that our senescent cell type was not a macrophage. Recent studies found that senescent Schwann cells can inhibit axonal growth after injury in aged mice [27, 28], so we next conducted additional mIHC stains in INK mice for p16 (senescent cells), SOX10 (pan Schwann cell marker) [29], and P0 (myelinating Schwann cells) [30]. Quantification of p16^+^ SOX10^+^ or p16^+^P0^+^ cells revealed no changes across groups indicating that senescent Schwann cells were not driving CIPN (**Extended Data Fig. 6A-D**).

To continue our investigation into the identity of the senescent cell type in an unbiased way, we turned to single cell RNA sequencing (scRNA-seq). Mice were treated with Veh or PTX according to our 9-day timeline and the hind paw skin was digested into a single cell suspension before sorting. Given our result that CD45^+^ cells do not play a role in our model of CIPN, CD45^-^ cells were collected and subject to downstream scRNA-seq analysis (**Extended Data Fig 7A-C**). 12 cell types were identified through known skin cell markers (**Extended Data Fig. 8**), including 4 fibroblast clusters, red blood cells (RBCs), smooth muscle cells, endothelial cells, sebaceous gland cells, keratinocytes, Schwann cells and 2 unknown cell clusters (**Fig. 4A**). To identify senescent cells, we looked for clusters expressing Cdkn2a, the gene encoding for p16. We found that Cdkn2a expression was restricted to the PDGFRα^+^ fibroblast clusters (**Fig. 4B**). Upon reclustering, we identified 7 distinct fibroblast clusters and found that Cdkn2a was substantially increased in cluster 7 in PTX treated mice (**Fig. 4C-D**). Cluster 7 also contained increased expression of senescent and SASP markers, including Cdkn1a (p21), Il6, Il1, and Mmp2 (**Fig. 4E-F**). To confirm the expression of p16 in this cluster (**Fig. 4G**), we conducted mIHC for p16 and PDGFRα^+^ in paw tissues from INK^+^ and INK^-^ mice treated with PTX. We found that PTX treatment increased p16^+^PDGFRα^+^ fibroblasts that were reduced with AP treatment (**Fig. 4H-I**). These results suggested that senescent PDGFRα^+^ fibroblasts located within the dermis of the skin drive axonal denervation and contributed to CIPN.

**Figure 4:**
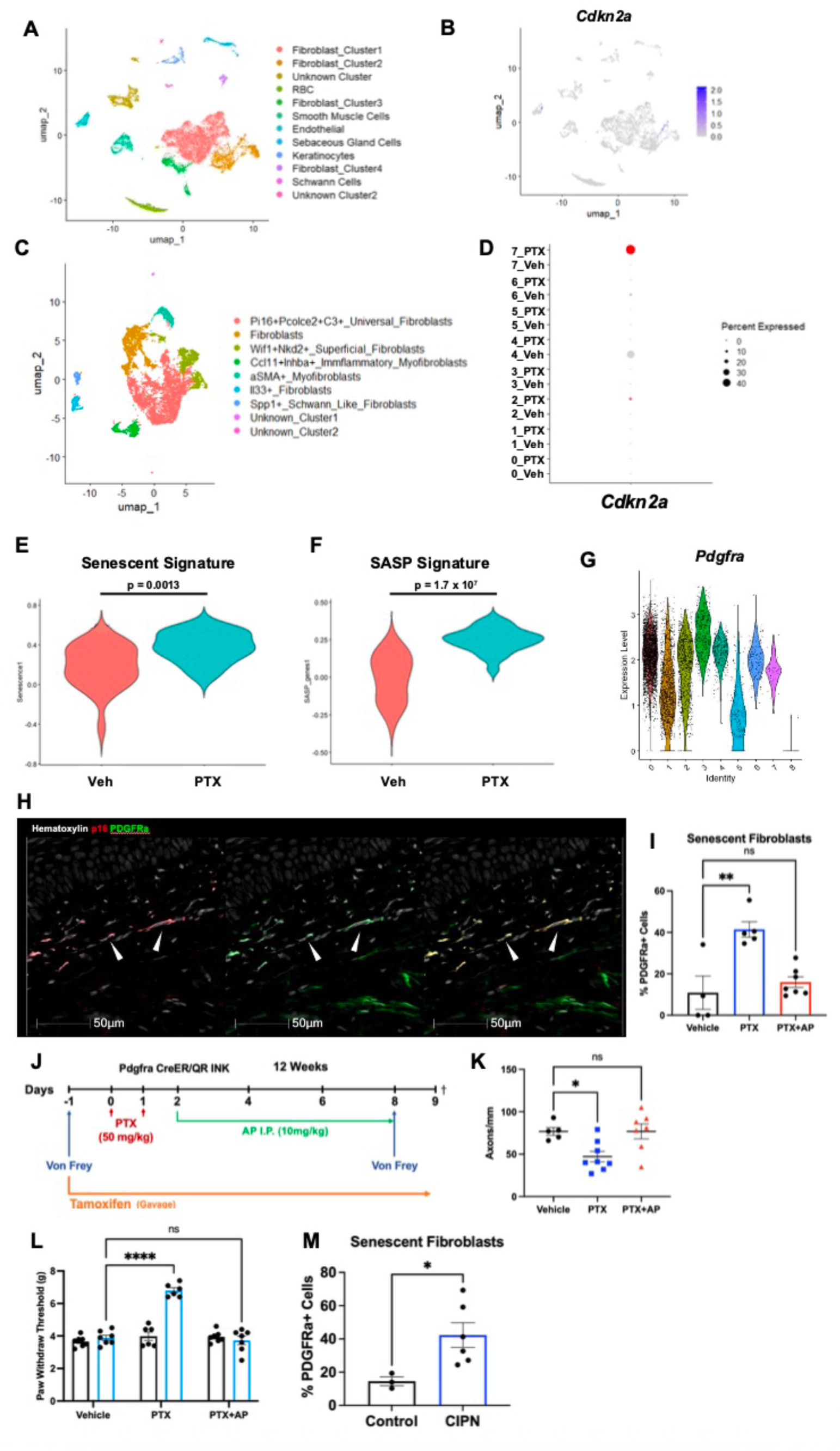
Dermal senescent fibroblasts drive CIPN. **(A**) UMAP of CD45^-^ cells from hind paw skin of 12-week-old mice treated with Veh or PTX (Pooled n=5). **(B)** UMAP of *Cdkn2a* (p16) expression across all cell type clusters. **(C)** UMAP of fibroblast clusters sub-setted from A. **(D)** Dot plot displaying gene expression parameters including percentage of cells (size of black dots) and expression levels (colored scale bar) of Cdkn2a (p16) within fibroblast clusters and either Veh or PTX treatment (y-axis). **(E)** Violin plot displaying the expression of senescence related genes in fibroblast cluster 7 of either Veh or PTX treated groups. **(F)** Violin plot displaying the expression of senescence related genes in fibroblast cluster 7 of either Veh or PTX treated groups. **(G)** Violin plot of Pdgfrα expression across all fibroblast clusters. **(H)** mIHC were performed on hind paw samples of 12-week-old INK-ATTAC mice using p16 (red) and PDGFRa (green) specific antibodies and nuclei were counterstained with hematoxylin and shown as white. Cells double positive for p16 and PDGFRα are denoted by white arrowheads. Individual stains were pseudo colored as indicated and then merged (last panel). **(I)** Quantification of senescent fibroblasts (p16+ PDGFRα^+^) out of total fibroblasts (PDGFRα^+^). Veh (n=4), PTX (n=5), PTX+AP (n=7). One-way ANOVA was performed, data are represented as mean ± SEM. **p< 0.01; ns, not significant. **(J)** Schema: Tamoxifen and AP delivery for PDGFRα CreER QR mice that received either Veh or PTX. **(K)** Quantification of innervating axons from PDGFRa CreER QR mice; (One-way ANOVA was performed, data are represented as mean ± SEM. *p< 0.05; ns, not significant). **(L)** Von Frey results on Day -1 (black) and Day 8 (blue) from same mice shown in K; (2-way ANOVA was performed, data are represented as mean ± SEM. ****p <0.001; ns, not significant). **(M)** Quantification of IHC for p16^+^ area our of total tissue area from same skin punch biopsies from either Control (n=3) or CIPN (n=6) patients. One-tailed Student t-test was performed, data are represented as mean ± SEM. *p< 0.05.

### Elimination of senescent fibroblasts prevents CIPN

To confirm a role for senescent PDGFRα^+^ fibroblasts in CIPN, we developed a new mouse model that expressed a lox-stop-lox (LSL) INK-ATTAC gene cassette (QR mouse) (**Extended Data Fig 9A**) [31]. To characterize the QR mouse, we crossed it to the Ella-Cre (ubiquitously expressed, JAX # 003724) mice to remove the LSL systemically so that we could detect expression in any cell undergoing senescence (**Extended Data Fig. 9B**). QR^+/+^ mice were treated with PTX resulting in increased p16^+^ that was reduced upon AP treatment (**Extended Data Fig. 9C**), indicating that the transgene is active. Importantly, AP prevented the development of numbness in mice treated with PTX (**Extended Data Fig. 9D**), indicating the suicide gene in the QR mouse recapitulates our original findings in the INK^+^ mice.

Having established the QR mouse effectively eliminates senescent cells, we next wanted to restrict the suicide gene expression to PDGFRα^+^ fibroblasts. Thus, we mated the QR mouse to the PDGFRα^+^-CreER^T2^ mouse (JAX # 018280) to generate PDGFRα^+^-CreER;QR (PDGFRα^+^-QR) mice. To activate Cre in PDGFRα^+^ fibroblasts, Cre^+^ and Cre^-^ mice were gavaged with 75mg/kg Tamoxifen (TAM) every other day starting one day before the first PTX dose and continued throughout the remainder of the experiment. Mice were also treated with AP (10mg/kg) every other day beginning after their second PTX dose. Mice underwent Von Frey assessment before PTX treatment and at endpoint (**Fig. 4J**). We found that Cre^-^ mice treated with PTX alone or PTX + AP displayed denervation and increases in PWTs, while Cre^+^ mice treated with PTX + AP displayed similar innervation and PWT as Veh treated animals, indicating that PDGFRα^+^ cells drove axon denervation (**Fig. 4K-L**).

In a complementary approach, we found that fibroblast cluster 7 also displayed increased expression of the fibroblast marker Acta2 (encoding for αSMA) (**Extended Data Fig. 10A**). Therefore, we carried out mIHC for αSMA and p16 in INK paw tissues. We observed that PTX increased αSMA^+^p16^+^ cells that were decreased by treatment with AP (**Extended Data Fig 10B-D**). Thus, we crossed the QR mouse with the Acta2-CreER^T2^ mouse (JAX #032758) to generate αSMA-CreER QR (αSMA-QR) mice. αSMA-Cre^+^ and αSMA-Cre^-^ mice were given chow compounded with TAM *ad libitum* starting one day before the first PTX dose and continued treatment throughout the remainder of the experiment. Mice were also treated with AP (10mg/kg) every other day beginning after their second PTX dose. Mice underwent Von Frey assessment before PTX treatment and at endpoint (**Extended Data Fig. 10E**). As expected, we found that αSMA-Cre^-^ mice treated with PTX alone or PTX + AP displayed increased PWTs, while αSMA-Cre^+^ mice treated with PTX + AP displayed similar PWTs to Veh treated animals (**Extended Data Fig. 10F**). Use of these two-cell type specific mouse models established that senescent fibroblasts within the dermis of the skin drive CIPN.

### Senescent fibroblasts are present in the skin of patients with CIPN

Since senescent fibroblasts were the drivers to our phenotype in murine models, we wanted to determine if these cells were present in patients with CIPN. To answer this, we returned to our skin punch biopsies from patients with CIPN or age matched controls [21] and conducted mIHC for p16 and PDGFRα. Quantitation of dual positive cells (p16^+^PDGFRα^+^) showed an increase in senescent fibroblasts in patients with CIPN compared to healthy controls (**Extended Data Fig. 11A-B**) (**Fig. 4M**). These results illustrate that chemotherapy-induced senescent skin fibroblasts are present in patients experiencing CIPN.

### MK2 pathway mediated SASP drives CIPN

The SASP can elicit a wide array of tissue responses [9] and our finding that senescent fibroblasts operated upstream of neuronal SARM1 activation raised the possibility that SASP drives CIPN. Previously we demonstrated that p38MAPK-MK2 pathway activation stabilizes SASP mRNAs, increasing protein expression [32, 33]. Further, we showed that we could rescue senescence-induced bone loss by treating mice with ATI450, an MK2 inhibitor to limit SASP expression [34] [35]. These findings led us to ask if targeting MK2 could prevent CIPN. To address this, we first treated 12-week-old MK2 knockout mice with Veh or PTX (**Fig. 5A**). As expected, IHC staining for p16 revealed that p16^+^ senescent cells were retained in MK2 KO mice, but p16^+^ cells showed reduced levels of phosphorylated HSP27 (pHSP27), a downstream target of MK2, in PTX treated MK2 KO mice (**Fig. 5B-C**) (**Extended Data Fig.12A-B**). Despite the presence of senescent cells and in contrast to WT mice (**Fig. 1**), MK2 KO mice did not experience axonal loss (**Fig. 5D-E**) and failed to develop symptoms (**Fig. 5F**), demonstrating that MK2 signaling contributes to CIPN.

**Figure 5:**
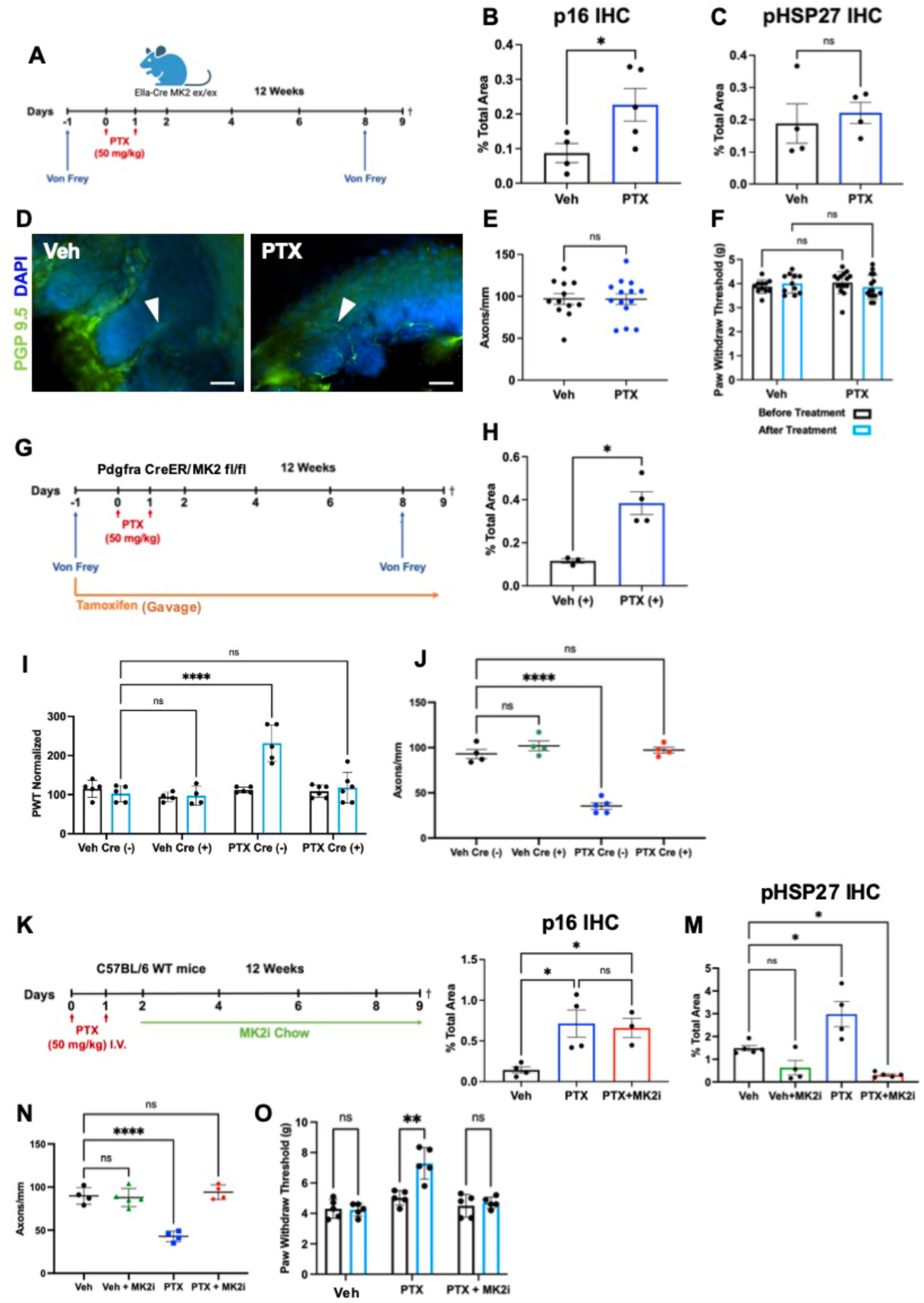
MK2 pathway mediated SASP prevents CIPN. **(A**) Schema: Veh or PTX treatment delivery for MK2 KO mice treated at 12-weeks-old. **(B)** Quantification of IHC for p16^+^ area our of total tissue area from same mice shown in E. Two-tail student t-test was performed, data are represented as mean ± SEM. *p< 0.05. **(C)** Quantification of IHC for pHSP27^+^ area our of total tissue area from same mice shown in E. Two-tail student t-test was performed, data are represented as mean ± SEM. *p< 0.05. **(D)** Representative IF images for PGP9.5 (green) and DAPI (blue) in hind paw sections from 12-week-old MK2 KO mice treated with Veh (n=12) or PTX (n=14). White arrowheads indicate innervating axons. **(E)** Quantification of innervating axons from same mice shown in B; (Two-tail student t-test was performed, data are represented as mean ± SEM. ns, not significant). **(F)** Von Frey results on Day -1 (black) and Day 8 (blue) from same mice shown in B; (2-way ANOVA was performed, data are represented as mean ± SEM; ns, not significant). **(G)** Schema: Tamoxifen delivery for PDGFRα CreER MK2 fl/fl mice that received either Veh or PTX. **(H)** Quantification of IHC for p16^+^ area out of total tissue area from Cre^+^ PDGFRα CreER MK2 fl/fl mice Veh Cre^+^ (n=3), PTX Cre^+^ (n=4). Two-tailed Student t-test was performed, data are represented as mean ± SEM. *p< 0.05. **(I)** Von Frey results on Day -1 (black) and Day 9 (blue) from Cre^-^ and Cre^+^ PDGFRα CreER MK2 fl/fl mice treated with Veh or PTX. Veh Cre^-^ (n=4), Veh Cre^+^ (n=4), PTX Cre^-^ (n=5) and PTX Cre^+^ (n=6). 2-way ANOVA was performed, data are represented as mean ± SEM. ****p< 0.001; ns, not significant. **(J)** Quantification of innervating axons from PDGFRα CreER MK2 fl/fl mice; Veh Cre^-^ (n=4), Veh Cre^+^ (n=4), PTX Cre^-^ (n=5) and PTX Cre^+^ (n=5) One-way ANOVA was performed, data are represented as mean ± SEM. ****p< 0.001; ns, not significant. **(K**) Schema: MK2i treatment schedule for 12-week-old WT mice treated with Veh or PTX. CDD2231 (MK2i) was used for these studies. **(L)** Quantification of IHC for p16^+^ area out of total tissue area from MK2i prevention study. Veh (n=4), PTX (n=4), and PTX+MK2i (n=3). One-way ANOVA was performed, data are represented as mean ± SEM. *p< 0.05. **(M)** Quantification of IHC for pHSP27^+^ area out of total tissue area from same mice MK2i prevention study. Veh (n=4), Veh+MK2i (n=4), PTX (n=4), and PTX+MK2i (n=3). One-way ANOVA was performed, data are represented as mean ± SEM. *p< 0.05. **(N)** Quantification of innervating axons from MK2i experiment; Veh (n=4), Veh+MK2i (n=5), PTX (n=4), PTX+MK2i (n=4). One-way ANOVA was performed, data are represented as mean ± SEM. ****p< 0.001; ns, not significant. **(O)** Von Frey results on Day -1 (black) and Day 8 (blue) from same mice shown in N; (2-way ANOVA was performed, data are represented as mean ± SEM. **p <0.01; ns, not significant).

To ask if MK2 activity in senescent fibroblasts was sufficient to induce CIPN, we crossed the B6N.Cg-Mapkapk2^tm2.1Yaff^/J (MK2 floxed) mouse (JAX # 032450) with the PDGFRα^+^-CreER^T2^ mouse to generate PDGFRα^+^-CreER MK2 fl/fl (PDGFRα^+^-MK2 KO) mice. Cre^-^ and Cre^+^ mice were gavaged with 75mg/kg Tamoxifen (TAM) every other day starting five days before the first PTX dose and continued treatment throughout the remainder of the experiment to remove MK2 from fibroblasts (**Fig. 5G**). To assess the induction of senescence, tissue sections from Veh and PTX treated mice were stained for p16. As expected, PTX treatment induced senescence in the skin as evidenced by p16 expression (**Fig. 5H**). To assess the impact of fibroblast-specific MK2 loss on the induction of CIPN, mice underwent baseline and endpoint Von Frey assessment. Cre^-^ mice treated with PTX exhibited increased PWTs, while Cre^+^ mice displayed readings similar to both Veh groups (**Fig. 5I**). Cre^+^ PTX treated animals also maintained innervation levels similar to Veh controls while Cre^-^ PTX mice experienced denervation (**Fig. 5J**). These results indicate that MK2 pathway activation in senescent fibroblasts is sufficient to drive CIPN.

Currently there are ongoing clinical trials using ATI450, a drug that inhibits the MK2 pathway in breast and pancreatic cancers [36], raising the prospect that MK2 inhibition (MK2i) could be leveraged in the clinic to prevent CIPN. Thus, we next asked if this inhibitor would be effective in preventing CIPN. Bolstered by our genetic data, we next treated 12-week-old WT C57Bl/6 mice with MK2i via compounded chow provided *ad libitum* (**Fig. 5K**). Consistent with the MK2 KO experiments, p16 expression was increased in PTX+MK2i mice, but pHSP27 was reduced when compared to PTX treatment alone mice (**Fig. 5L-M**) (**Extended Data Fig. 12C-D**), indicating that MK2i effectively limited pathway activation. Despite the activation of senescence in MK2i treated mice, PTX treated mice receiving MK2i failed to display axon denervation (**Fig. 5N**) or symptoms of CIPN (**Fig. 5O**). These exciting findings suggest that MK2i could prevent CIPN in patients.

### Eliminating senescent cells rescues CIPN

Senescent fibroblasts induce CIPN but whether their continued presence prevents re-innervation was an open question. This raised the possibility that if we could eliminate senescent fibroblasts in animals with established CIPN, we might see re-innervation and the reversal of CIPN [37]. To address this, we next treated mice with PTX and allowed them to develop CIPN as demonstrated by an increase in PWT on day 9 (**Fig. 6A-B**). PTX treated mice with established CIPN were then treated with AP every other day for 18 days and subject to a third Von Frey assay. At endpoint, PTX treated animals displayed an increased number of p16^+^ cells that was reduced by AP treatment (**Fig. 6C**) (**Extended Data Fig. 13A**), indicating that senescent cells persist long after PTX treatment. Excitingly, AP treatment of mice with established CIPN returned their PWT to pretreatment levels whereas mice treated with PTX displayed a reduced PWT relative to pretreatment, indicating that they had progressed to pain (**Fig. 6B**). As expected, PTX treated mice displayed less intervention compared to Veh treated mice or mice treated with PTX+AP (**Fig. 6D**). Together these data demonstrate that the elimination of senescent fibroblasts rescues established CIPN and raises the possibility that patients with established CIPN could be treated.

**Figure 6:**
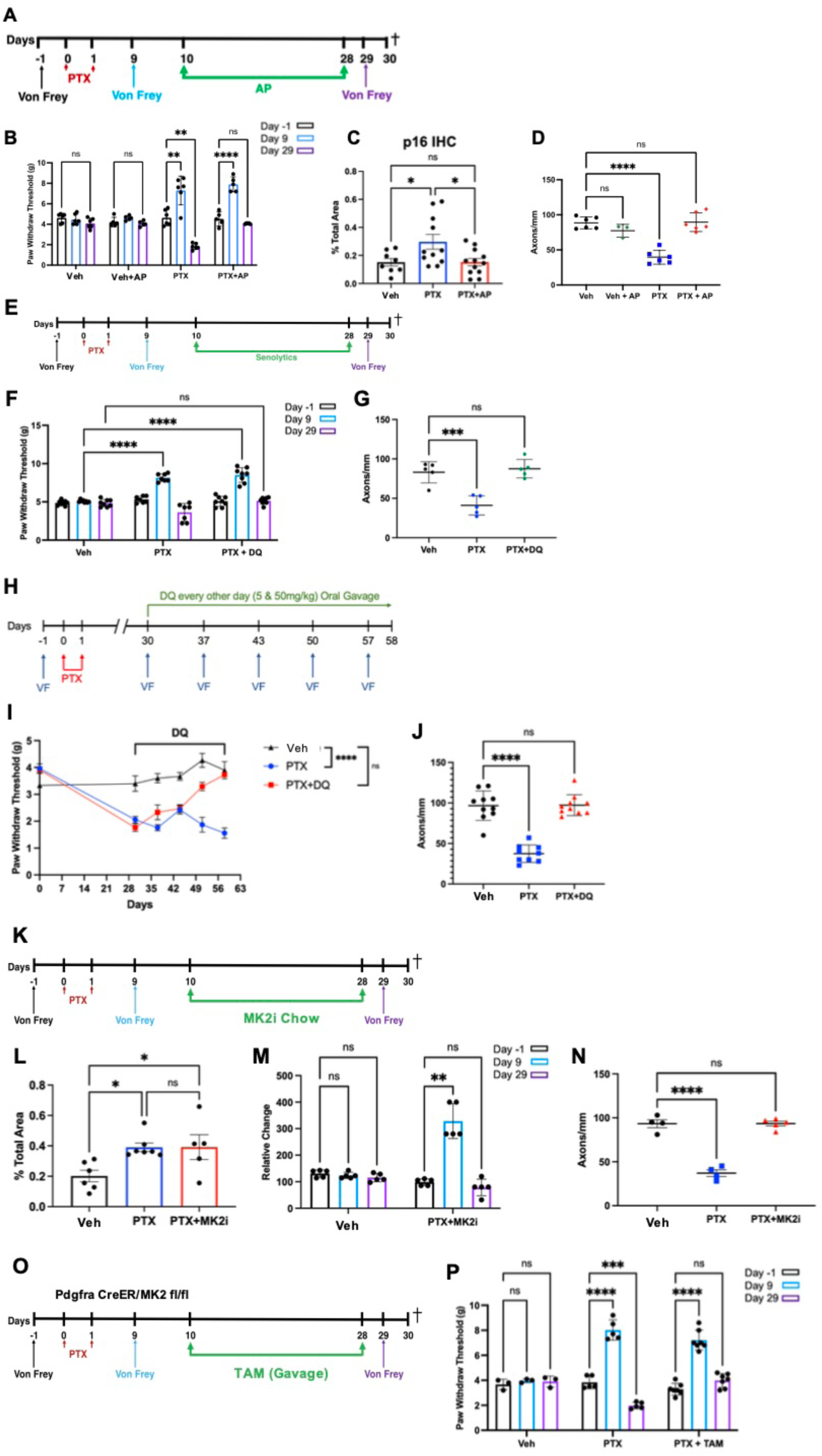
Genetic or pharmacologic elimination of senescent cells rescues CIPN. **(A)** Schema: PTX and AP treatment regimen for INK-ATTAC rescue studies. **(B)** Von Frey results on Day -1 (black), Day 8 (blue), and Day 29 (purple) from same mice shown in D; (2-way ANOVA was performed, data are represented as mean ± SEM. **p<0.01; ****p <0.001; ns, not significant). **(C)** Quantification of p16^+^ area out of total tissue area. Veh (n=8), PTX (n=11), PTX+AP (n=12); (One-way ANOVA was performed, data are represented as mean ± SEM. *p< 0.05; ns, not significant). **(D)** Quantification of PGP9.5 innervating axons from same mice shown in B. Veh (n=6), Veh+AP (n=3), PTX (n=6), PTX+AP (n=6); (One-way ANOVA was performed, data are represented as mean ± SEM. ****p< 0.0001; ns, not significant). **(E)** Schema: PTX and DQ treatment regimen for senolytic rescue studies. **(F)** Von Frey results on Day -1 (black), Day 8 (blue), and Day 29 (purple) from same mice shown in G; (2-way ANOVA was performed, data are represented as mean ± SEM. ****p <0.001; ns, not significant). **(G)** Quantification of innervating axons from senolytic rescue studies. Veh (n=5), PTX (n=5, and PTX+DQ (n=5). One-way ANOVA was performed, data are represented as mean ± SEM. ***p< 0.001; ns, not significant. **(H)** Schema: PTX and DQ treatment regimen for pain rescue studies. **(I)** Von Frey results from pain rescue mice. Veh (black), PTX (blue), and PTX+DQ (red) from same mice shown in J; (2-way ANOVA was performed, data are represented as mean ± SEM. ****p <0.001; ns, not significant). **(J)** Quantification of innervating axons from pain rescue mice shown in H; Veh (n=10), PTX (n=10), PTX+DQ (n=10). (One-way ANOVA was performed, data are represented as mean ± SEM. ***p< 0.001; ns, not significant). **(K**) Schema: MK2i treatment schedule for 12-week-old WT mice treated with Veh or PTX. ATI450 (MK2i) was used for these studies. **(L)** Quantification of IHC for p16^+^ area out of total tissue area from MK2i rescue studies. One-way ANOVA was performed, data are represented as mean ± SEM. *p< 0.05. **(M)** Von Frey results on Day -1 (black), Day 8 (blue) and Day 29 (purple) from same mice shown in D; (2-way ANOVA was performed, data are represented as mean ± SEM. **p <0.01; ns, not significant). **(N)** Quantification of innervating axons from MK2i experiment; Veh (n=4), Veh+MK2i (n=5), PTX (n=4), PTX+MK2i (n=4). One-way ANOVA was performed, data are represented as mean ± SEM. ****p< 0.001; ns, not significant. **(O**) Schema: Tamoxifen delivery for PDGFRα CreER MK2 fl/fl mice that received either Veh or PTX for rescue studies. **(P)** Von Frey results on Day -1 (black), Day 8 (blue) and Day 29 (purple) from same mice shown in E; Veh (n=3), PTX (n=5), and PTX+TAM (n=5). 2-way ANOVA was performed, data are represented as mean ± SEM. ***p <0.001, ****p< 0.0001; ns, not significant.

Keeping patients in mind, we next asked if senolytics could rescue CIPN after it had developed. Mice were treated with PTX and allowed to develop CIPN as demonstrated by an increase in PWT on day 9. PTX treated mice that had developed CIPN were then treated with Veh or DQ for 18 days and subject to a third Von Frey assay (**Fig. 6E**). DQ treatment returned the PWT to pretreatment levels whereas mice treated with PTX plus Veh displayed a reduced PWT relative to pretreatment, indicative of pain (**Fig. 6F**). In agreement with the behavior test, PTX treated mice displayed less intervention compared to Veh treated mice or mice treated with PTX and DQ (**Fig. 6G**). Given that many patients progress from numbness to pain after completion of their chemotherapy cycles [38], we next wanted to ask if senolytics could rescue pain symptoms of CIPN. To address this, mice were treated with PTX and allowed to develop pain symptoms of CIPN as demonstrated by a decrease in PWT on day 30. PTX treated mice that had developed CIPN were then treated with DQ every other day for the next 5 weeks receiving a Von Frey assessment every week for signs of symptom improvement (**Fig. 6H**). After 5 weeks of DQ treatment, mice that had developed CIPN returned to PWTs similar to Veh controls. PTX only treated animals continued to exhibit pain like phenotypes (**Fig. 6I**). Axonal staining at endpoint also showed a rescue of denervation in PTX+DQ animals compared to PTX alone (**Fig. 6J**). The ability of senolytics like DQ to rescue CIPN in both phenotypes of numbness and pain strongly supports a role for senescent killing drugs in treating patients with CIPN.

### Therapeutic targeting of MK2 can rescue CIPN

Encouraged by our finding that MK2i could prevent CIPN and that senolytics could rescue established CIPN, we next asked how MK2i impacted established CIPN. To address this possibility, WT mice were treated with PTX and allowed to develop CIPN as demonstrated by an increase in PWT on day 9 (**Fig. 6K**). PTX treated mice that had developed CIPN were then placed on MK2i compounded chow for 18 days and subject to a third Von Frey assay. As expected, IHC for p16^+^ cells showed an increase in PTX+MK2i treated animals (**Fig. 6L**), confirming what was observed in our genetic INK rescue studies (**Fig. 6B**). MK2i treatment returned the PWT to pretreatment levels whereas mice treated with PTX plus Veh displayed a reduced PWT relative to pretreatment, indicative of pain (**Fig. 6MC**). In agreement with the Von Frey, PTX treated mice displayed less intervention compared to Veh treated mice or mice treated with PTX and MK2i (**Fig. 6N**). These studies indicate that MK2 mediated SASP drives and maintains CIPN.

We next wanted to ask if CIPN could be rescued by limiting SASP expression in fibroblasts. To do this, PDGFRα^+^-MK2 KO Cre^+^ mice were treated with PTX and allowed to develop CIPN as demonstrated by an increase in PWT on day 9 (**Fig. 6P**). PTX treated mice that had developed CIPN were then gavaged with 75mg/kg TAM every other day for 18 days to activate Cre and remove MK2 and subject to a third Von Frey assay (**Fig. 6O**). Cre^+^ mice treated with PTX exhibited a PWT similar to pretreatment levels whereas Cre^-^ mice treated with PTX displayed reduced PWT relative to pretreatment, indicative of pain (**Fig. 6P**). These results indicate that the loss of MK2 specifically in senescent fibroblasts is sufficient to rescue existing CIPN.

## DISCUSSION

Chemotherapy-induced peripheral neuropathy (CIPN) represents a major clinical challenge, frequently forcing dose de-escalation or the cessation of essential cancer therapies, thereby directly compromising patient survival [1]. Further, for those who manage to beat their cancer, CIPN can linger for a lifetime where it negatively impacts quality of life. For decades, CIPN has been thought of as a cell-autonomous axonal vulnerability, leading researchers to focus on how direct neurotoxic insults result in CIPN [39]. Here, we present a fundamental shift in thinking about what drives CIPN. Indeed, we show that CIPN is driven by a non-neuronal, stromal-derived cell in the distal skin. By deploying single-cell transcriptomics, cell-type-specific inducible Cre models, and pharmacological interventions, we demonstrate that paclitaxel and cisplatin induce a distinct population of senescent dermal fibroblasts. These cells dictate both the initiation and long-term maintenance of peripheral neuropathy through an MK2-dependent SASP that acts upstream of neuronal SARM1.

The localization of the senescent fibroblasts provides insight into peripheral neuropathies. While previous *in vitro* studies suggested that chemotherapeutics induce senescence within the DRG [18], our *in vivo* analyses reveal that the senescent cells are restricted to the distal dermis. This anatomical localization suggests that the distal axons are extremely responsive to alterations in their immediate microenvironment and that these changes can initiate axonal withdrawal. Further, our rescue studies suggest that the continued presence of senescent fibroblasts precludes reinnervation of the peripheral tissue but upon elimination of senescent fibroblasts, the axons can rapidly reinnervate the tissue and reverse the symptoms of CIPN.

Our findings also suggest that senescence functions upstream of neuronal SARM activation. By characterizing SARM1-deficient mice that are protected from CIPN [24], we find that upon paclitaxel treatment mice still accumulate senescent cells in the skin yet fail to develop CIPN. In an effort to understand how senescent fibroblasts stimulate SARM1, we turned to the SASP, which is known to have pleotropic effects on surrounding tissue [40] and consists of numerous secreted proteins including IL6, TNFα, and HMGB1, all of which have been implicated in CIPN [10, 11, 41]. Importantly, our data mechanistically reveal the central role the p38MAPK-MK2 pathway plays in CIPN. Deletion or inhibition of MK2 selectively halts the stabilization of SASP transcripts (such as *Il6*, *Il1a*, *HMGB1,* and *Mmp2,* etc.) without preventing the induction of senescence. Consequently, silencing this single node within the dermal stroma is sufficient to preserve peripheral innervation and prevent sensory dysfunction.

From a translational perspective, the most compelling aspect of this study is the therapeutic opportunities it reveals; indeed, our work shows that CIPN can be prevented and even reserved by senolytics or an MK2 inhibitor. In the clinic, many patients develop symptoms ranging from acute numbness to chronic, debilitating neuropathic pain after completing their chemotherapy cycles [4]. Our data shows that senescent dermal fibroblasts persist in the skin post-chemotherapy, acting as a continuous source of neurotoxic SASP that block re-innervation. By eliminating these cells or by halting their secretory activity (via the MK2 pathway inhibitor ATI-450) weeks after the establishment of CIPN, we achieved a striking reversal of both tactile numbness and late-stage neuropathic pain. This therapeutic reversal directly coincided with the regeneration of peripheral axons into the epidermis. Thus, deploying an MK2 inhibitor alongside frontline chemotherapies could concurrently suppress primary tumor progression and safeguard patients from the devastating, treatment-limiting toxicities of peripheral neuropathy.

## MATERIALS and METHODS

### Animal studies

All mice were housed and all animal procedures were approved by Washington University’s Institutional Animal Care and Use Committee (IACUC). Mice were maintained on a 12-hour light/12-hour dark cycle, with ambient temperature between 20–22.2 °C and relative humidity of 30–70%. Mice were maintained on a regular chow diet (PicoLab Rodent Diet 20, Cat. No. 5053) formulated with 20% protein. C57BL/6 J (JAX, #000664) mice were purchased from the Jackson Laboratory. SARM1 KO (JAX, #018069) mice were purchased from the Jackson Laboratory. The C57BL/6J INK mouse line was a kind gift from Dr. Darren Baker. QR^+/+^ mice were generated by crossing the QR mouse with B6.FVB-Tg(EIIa-cre)C5379Lmgd/J (JAX # 003724) mice purchased from the Jackson Laboratory. To generate the PDGFRα^+^-CreER;QR (PDGFRα^+^-QR) mice, QR mice were crossed with B6N.Cg-Tg(Pdgfra-cre/ERT)467Dbe/J (JAX # 018280) mice purchased from the Jackson Laboratory. Mice were gavaged with tamoxifen (Cayman Chemical Company #13258) dissolved in corn oil (ThermoFisher 405430025) at 75mg/kg every other day, starting 1 day before the first dose of PTX. To generate αSMA-CreER QR (αSMA-QR) mice, QR mice were crossed with B6(Cg)-Tg(Acta2-cre/ERT2)1Ikal/J (JAX # 032758) mice purchased from Jackson Laboratory. Mice were given diet gel (Clear H_2_O Diet Gel Recovery Cat #:72-06-5022) compounded with 60mg of tamoxifen (Cayman Chemical Company #13258) *ad libitum* starting one day before the first PTX dose and continued treatment throughout the remainder of the experiment. to generate PDGFRα^+^-CreER MK2 fl/fl (PDGFRα^+^-MK2 KO) mice, B6N.Cg-Tg(Pdgfrα-cre/ERT)467Dbe/J (JAX # 018280) were crossed with B6N.Cg-Mapkapk2^tm2.1Yaff^/J (JAX #032450) mice, both purchased from Jackson Laboratory. Mice used in this study were genotyped by TransnetYX (an automated genotyping company, USA).

### Drug Treatments

Paclitaxel (TOCRIS, a Biotechne brand) stock solution was prepared in a 1:1 solution of 100% ethanol and Kolliphor. Stock solution was further diluted in PBS prior to injection resulting in a final concentration of 7.5mg/mL. PTX was administered in two doses at 50mg/kg via tail vein (IV). Cisplatin (Enzo Cat #: ALX-400-040-M050) was dissolved into saline (Hospira NDC 0409-4888-06) and injected intraperitoneal (i.p.) at 2.3 mg/kg for 2 cycles, with 5 consecutive daily injections in each cycle and a 5-day rest between the two cisplatin treatments. AP20187 (Batch No.: A113-20, Chemvada Life Sciences, San Diego, CA) and prepared with 100% ethanol: polyethylene glycol 400: 2% Tween-20 in molecular water at 4:10:86. AP20187 was administered for a week at 10mg/kg thrice through i.p. injections.

### MFP Tumor Injections

PyMT-Bo1 were cultured at 37°C with 5% CO_2_ in complete media DMEM supplemented with 10% heat-inactivated FBS, 100 μg/ml streptomycin, 100 IU/ml penicillin, and 1 mM sodium pyruvate. All cell lines were tested for Mycoplasma. Tumor cells (1 × 10^5^) resuspended in a 1:1 PBS/Matrigel mixture (Corning, 354234; 50 μL) were implanted orthotopically into the mammary fat pad (MFP) of 12-week-old female mice. Tumor growth was monitored with caliper measurements every other day starting 7 days after implantation, and tumor volume was calculated using the formula V = 0.5 × (length × width^2^).

### Oral feeding of MK2i compounded chow

The MK2 small-molecule inhibitor CDD2231 (MK2i) (Aclaris Therapeutics, Inc.) was compounded at 1000 PPM. CDD2231 was compounded into Research Diets Inc., catalog number 5001. All C57BL/6 mice were fed *ad libitum*. Mice were randomized onto inhibitor-containing or regular chow one day after their last dose of PTX and remained on the chow until day of sacrifice for prevention studies. For rescue studies the MK2 small inhibitor ATI450 (MK2i) (Aclaris Therapeutics, Inc.) was compounded at 1000 PPM. ATI450 was compounded into Research Diets Inc., catalog number 5001.For these studies, mice were placed on chow on day 9 after confirmation of numbness and remained on chow for 21 days until day of sacrifice. All C57BL/6 mice were fed *ad libitum*.

### Behavioral studies

For prevention behavioral studies, all mice were tested 1 day before PTX treatment and 1 day before date of sacrifice. For rescue behavioral studies, all mice were tested 1 day before PTX treatment, 9 days after first PTX treatment, and 28 days after first PTX treatment. The experimenters were blind to the genotype.

### Von Frey

Mice were habituated individually in 10 × 10 × 15 cm open Plexiglass boxes with perforated lids on wire mesh for 30 minutes. After mice stopped exploring and were resting, von Frey testing was performed according to the up-down method [42]. Briefly, the middle of the plantar surface of the hind paw between the footpads was stimulated for 2 s with a filament connected to the BioSeb Electronic von Frey (BIO-EVF-WRS). This was repeated three more times. Right and left hind paws were tested separately. Von Frey experiments in the beginning and at the end of the experiment were done at the same time of the day.

### Immunohistochemistry (IHC) staining and quantification

Specific information on vendor, catalog number, and dilution ratio of any antibody used for IHC or IF staining is listed in Supplementary Table 1.

Harvested hind paws were fixed in either 10% normal buffer formalin over night at 4°C for 24–48 hours, washed in phosphate buffered saline (PBS), and preserved in 70% ethanol at 4 Celsius until being processed into formalin-fixed paraffin-embedded (FFPE) blocks. FFPE sections of 5–7 μm thickness were stained on a Leica BOND Automated IHC/ISH Stainer. For p16 staining, mouse tumor sections were first stained with anti-CDKN2A (pH 6.0 antigen retrieval) in combination with rabbit anti-rat antibody using the BOND Polymer Refine Detection System. Slides were mounted and scanned using a Zeiss Axio Scan.Z1 slide scanner, and the resulting images were analyzed using Indica Lab’s HALO computational pathology software.

mIHC staining on human or mouse tissue sections were performed on a Leica BOND Automated IHC/ISH Stainer using the BOND Polymer Refine Detection System in combination with goat anti-rabbit Fab fragment, rabbit anti-rat antibody, 3-Amino-9-ethylcarbazole (AEC) chromogenic substrate (Abcam ab64252) and hematoxylin (DAKO S3301). Human skin biospy tissue sections were first stained with anti-CDKN2A antibody (pH 6.0 antigen retrieval). After the staining was finished, slides were mounted with aqueous mounting media (Vector H-5501) and scanned. The slides were then submerged in TBS-T buffer at 4° Celsius overnight to remove the coverslip. After 3 washes in diH2O, the slides were incubated in 50% ethanol for 5 minutes and then transferred to acidified (1% HCl) 70% ethanol and incubated for 10 minutes with agitation to remove AEC chromogen and hematoxylin. The slides were then incubated with 100% ethanol for 15 minutes with agitation and was rehydrated and stored in TBS-T at 4° Celsius until next staining cycle. Such staining-scanning-stripping cycle was repeated for each of the following markers in the listed order: αSMA (pH 6.0 antigen retrieval), and PDGFRα (pH 8.0 antigen retrieval). Mouse paw tissues were processed using the staining-scanning-stripping procedures as described above and stained for the following markers in the listed order to probe senescent cell populations: CDKN2A (pH 6.0 antigen retrieval), MPZ (pH 6.0 antigen retrieval), αSMA (pH 6.0 antigen retrieval), and F4/80 (pH 6.0 antigen retrieval). Mouse paw tissues were processed using the staining-scanning-stripping procedures as described above and stained for the following markers in the listed order to probe additional senescent cell populations: CDKN2A (pH 6.0 antigen retrieval), SOX10 (pH 6.0 antigen retrieval), αSMA (pH 6.0 antigen retrieval), and PDGFRα (pH 8.0 antigen retrieval). All scanned bright field images for mIHC staining were deconvoluted into pseudo-fluorescent images and merged using Indica Lab’s HALO computational pathology software.

### Immunofluorescence PGP9.5 staining and axon quantification

Harvested hind paws were fixed in freshly prepared Zamboni’s fixative (0.12% picric acid, 2% paraformaldehyde in 0.1 M PBS, pH 7.4) over night at 4°C for 24–48 hours, washed in phosphate buffered saline (PBS), and immersed in 30% sucrose overnight at 4°C. They were then frozen in O.C.T. before 50 -μm thick cross-sections were cut at the cryostat (Leica CM1860). Series of sections were stored prior to processing at −20°C in a cryoprotectant comprising 30% sucrose and 33% ethylene glycol in PBS. A series of sections was thoroughly rinsed in 1x PBS, immersed in 10% normal donkey serum (NDS) and 0.1 % Triton™ X-100 in PBS (PBS-T) for 1 h, and transferred into rabbit anti-Protein Gene Product 9.5 (1:1000, EMD Millipore) overnight at 4°C. The next day, sections were thoroughly rinsed in PBS-T and placed in Alexa Fluor 594-conjugated donkey anti-rabbit secondary antibody (Invitrogen) at a dilution of 1:500 for 2 hours at room temperature. After further rinsing, sections were mounted on gelled slides and cover slipped using Vectashield with DAPI (Vector Laboratories) to allow visualization of nuclei.

The part of the plantar surface containing the footpad was identified and imaged on a Nikon Eclipse 90i microscope using a 40x oil objective and a Nikon Cool Snap HQ2 camera. Axons crossing into the epidermis over the entire medial lateral extension of the footpad were determined by examining the entirety of the 50 μm stack. Axons that crossed the basement membrane were counted, whereas secondary branching and epidermal nerve fragments that do not cross the basement were not [14]. Imaging and analysis were done with the samples blinded to the observer.

### SPiDER-**β**-gal staining

Mouse paws and DRG were fixed in freshly prepared Zamboni’s fixative (0.12% picric acid, 2% paraformaldehyde in 0.1 M PBS, pH 7.4) or 4% paraformaldehyde (PFA) overnight at 4°C, washed in phosphate buffered saline (PBS), and immersed in 30% sucrose overnight at 4°C. Tissues were then embedded in OCT compound (Fisher Health Care, Cat# 4585) and sectioned at 10 μm thickness using a cryotome (Leica CM1950). Cryosections were washed in PBS for 2 min and incubated with 5 μM SPiDER-β-gal probe (Dojindo Laboratories, Cat# SG02) at 37°C for 45 minutes in the dark. After staining, sections were washed with PBS and mounted using SlowFade Gold antifade reagent with DAPI (Invitrogen S36939) according to manufacturer instruction. Slides were scanned using the Zeiss Axio Scan Z1 fluorescence slide scanner and SPiDER-β-gal positive area was analyzed by HALO software.

### Bone marrow transplantation

Bone marrow transplantation was performed according to previously described protocols. Recipient C57BL/6 CD45.1 mice received two doses of 400 cGy 4 hours apart, followed by transplant of bone marrow by retro-orbital injection (i.v.). Irradiation was carried out using an X-ray irradiator (XRAD 320). Donor bone marrow was prepared from INK-ATTAC CD45.2 mice as follows: donor mice were sacrificed by CO_2_ inhalation, both femurs, tibias and ilia were extracted in a sterile setting and flushed using pulsed centrifugation to collect marrow. Bone marrow was reconstituted in cold sterile serum-free 1X HBSS and injected retro-orbital at a concentration of 5 million cells per 200 μl per mouse. Mice were monitored over 2 weeks for signs of radiation sickness. 6-week post irradiation, mice were given paclitaxel and sacrificed 9 days later.

### Single-cell RNA-sequencing

For scRNA-seq analysis of paws from 12-week-old WT mice treated with Veh or PTX, 5 biological replicates were pooled for each treatment group. Paws were collected from the hind paws of the mice and digested in DMEM F-12 media (Gibco 11320033) containing 2 mg/mL collagenase A (Millipore Sigma 10103578001), 1mg/mL Hyaluronidase (Cat #) and 2 U/mL DNase for 1 hour at 37° Celsius. Dissociated tissue suspension was then filtered through a 70 μm strainer and centrifuged at 1500 round per minute (rpm) for 5 minutes. Dissociated tissue was additionally filtered through a 40 μm strainer and centrifuged at 1500 round per minute (rpm) for 5 minutes. Cell pellets were then washed using FACS buffer (1% bovine serum albumin, 1 mM EDTA, 0.05% sodium azide in PBS) and incubated with Fc receptor blocking antibody for 10 minutes on ice. Cells were then stained for CD45. Stained samples were sorted on a Sony Synergy 5 laser, 22 colors cell sorter to separate CD45+ and CD45− cell populations. The collected samples were processed for library generation following the instructions of 10x Genomics Chromium X instrument (10x Genomic, Pleasanton CA). The quality of the prepared libraries was assessed and submitted for sequencing at the Genome Technology Access Center. Output from Cell Ranger Software was imported into R Studio and analyzed using Seurat [43]. For each Seurat object, genes expressed by less than 3 cells and cells expressing less than 1,000 or more than 8,000 genes were excluded. Cells with higher than 15% mitochondrial RNA content and cells with less than 100 or more than 250,000 counts were excluded as well. SCTransform was performed to normalize the expression matrix [44]. PCA and UMAP dimensional reduction was performed using the first 25 PCA components. We then conducted FindNeighbors and FindClusters functions to cluster cells. The FindAllMarkers function was used to identify signature genes for each cluster.

### Human CIPN Bulk RNA-sequencing analysis

For the analysis of the publicly available human skin biopsy bulk RNA-seq dataset (GEO: GSE228633), we utilized N=3 healthy and CIPN patient samples. Gene counts less than 10 were filtered out before DESeq2 analysis to identify differentially expressed genes.

## ACKNOWLEDGMENTS

We thank Dr. Katherine Weilbaecher and Dr. Valeria Cavalli for constructive input. Additionally, we thank Dr. Aaron DiAntonio and Lilliane Barber for assistance with in vitro systems and constructive input. We thank Gaurav Swarnkar for his assistance with the von Frey in the Musculoskeletal core, Washington University School of Medicine. We also thank Anupama Melam for assistance with Von Frey assays. We are also grateful for the core services provided by the Musculoskeletal Research Center (NIH P30-AR074992). This work was supported by NIH grants R01CA282810, R01AG093010, and R01AG088264 (S.A. Stewart), the U.S. Army Medical Research Acquisition Activity, 820 Chandler Street, Fort Detrick, MD 217025014, is the awarding and administrating acquisition office, and this was supported in part by the Office of the Assistant Secretary of Defense for Health Affairs, through the Breast Cancer Research Program, under award No. MBC181712. Opinions, interpretations, conclusions, and recommendations are those of the authors and are not necessarily endorsed by the Department of Defense. We also thank the Siteman Flow Cytometry & Fluorescence Activated Cell Sorting Pathology Core and Daniel Schweppe in the Siteman Flow Cytometry Core for help with sorting. In addition, we thank the Genome Technology Access Center (GTAC) in the Department of Genetics at Washington University School of Medicine for single cell RNA-sequencing. The Centers are partially supported by NCI Cancer Center Support Grant #P30 CA91842 to the Siteman Cancer Center and by ICTS/CTSA Grant #UL1 TR000448 from the National Center for Research Resources (NCRR), a component of the National Institutes of Health (NIH), and NIH Roadmap for Medical Research. This publication is solely the responsibility of the authors and does not necessarily represent the official view of NCRR or NIH. This work was also supported by the Siteman Cancer Center Investment Program (NCI Cancer Center Support Grant P30CA091842, Fashion Footwear Association of New York, and the Alvin J. Siteman Cancer Center, Siteman Investment Program (supported by The Foundation for Barnes-Jewish Hospital, Cancer Frontier Fund) to S.A. Stewart. The work was also supported by the Washington University Musculoskeletal Research Center (NIH T32 AR060719) and Molecular Oncology Training Program (NCI T32 CA113275) to T. Malachowski.

## Data Availability

The scRNA-seq datasets generated in this study are also available in the NCBI Gene Expression Omnibus (GEO) under accession number GSE342321 (samples derived from mice). Raw data can be found on Zenodo; 10.5281/zenodo.21723207.

## Author contributions

Conceptualization, data curation, formal analysis, investigation, methodology, visualization, writing–original draft, writing–review and editing, T.M. Investigation, S.M.B., and X.Y. Investigation and formal analysis, G.K.R., R.M., and D.D; Investigation and resources: Q.R., N.P.S., T.J.P., S.H. Conceptualization, resources, data curation, supervision, funding acquisition, writing– original draft, project administration, writing–review and editing, S.A.S.; all authors approved and provided comments on the submitted manuscript.

## DECLARATION OF INTERESTS

The authors declare no competing interests.

## Notes

### Competing Interest Statement

The authors have declared no competing interest.

