## Supplementary figures and images for "Senescent peripheral fibroblasts initiate chemotherapy-induced peripheral neuropathy"

### Ex.t Fig. 1

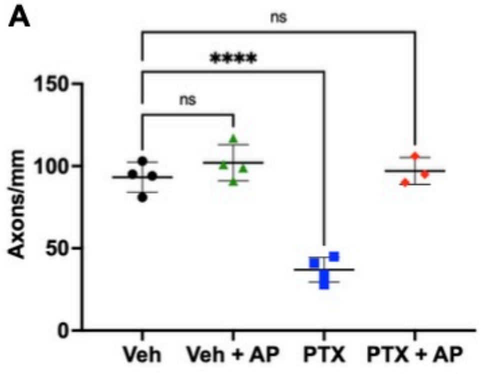


**Extended Data Fig. 1:**

### Ex.t Fig. 2

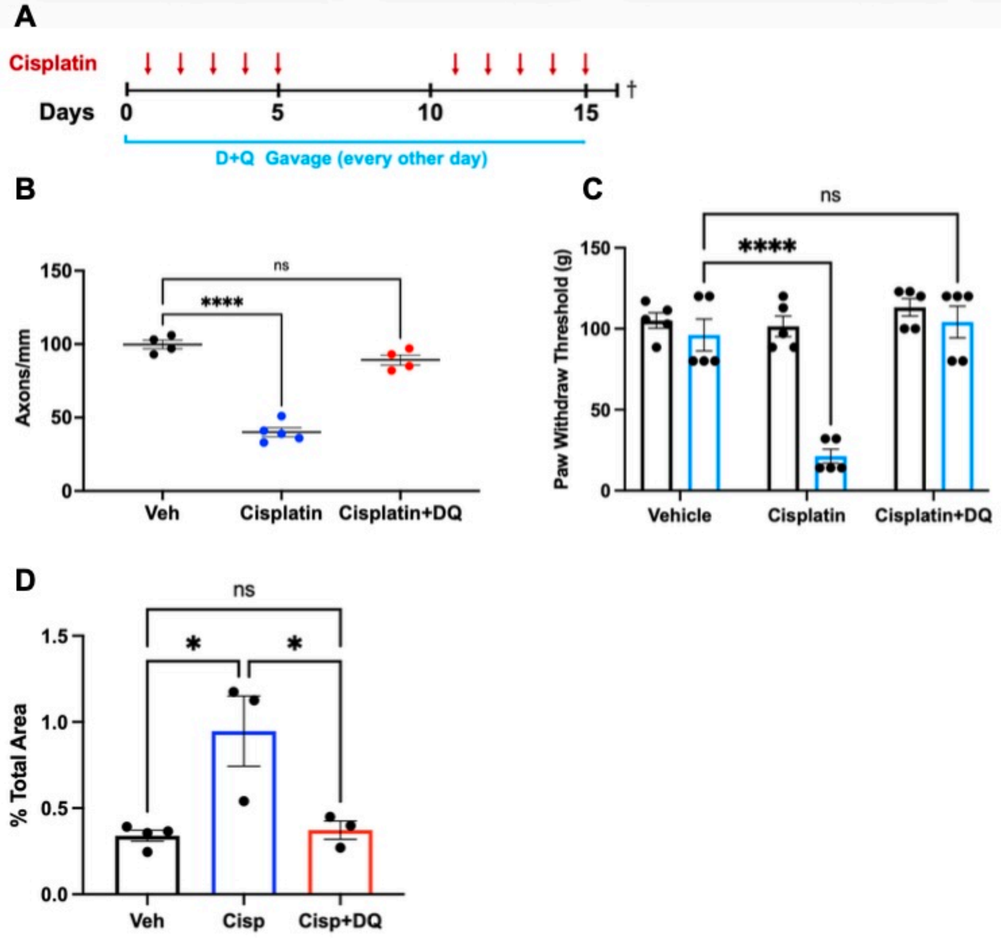


**Extended Data Fig. 2:**

### Ex.t Fig. 3

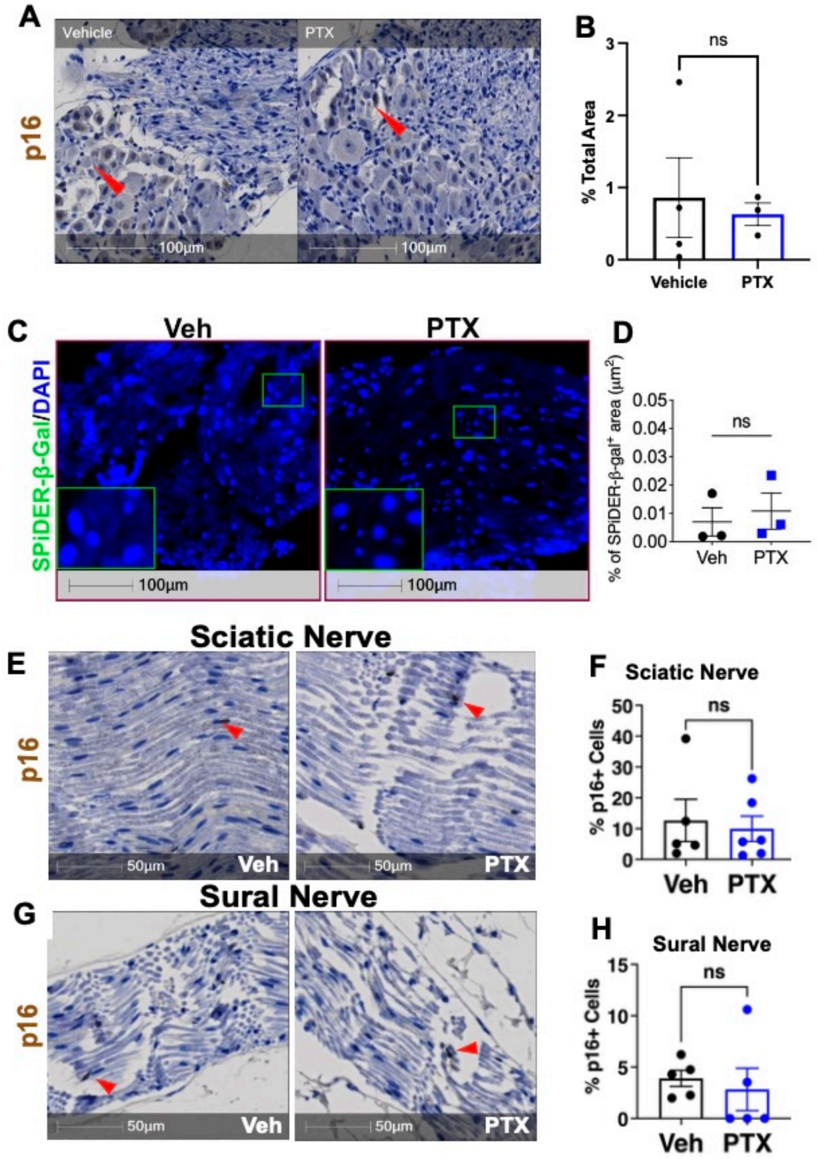


#### Extended Data Fig. 3:

### Ex.t Fig. 4

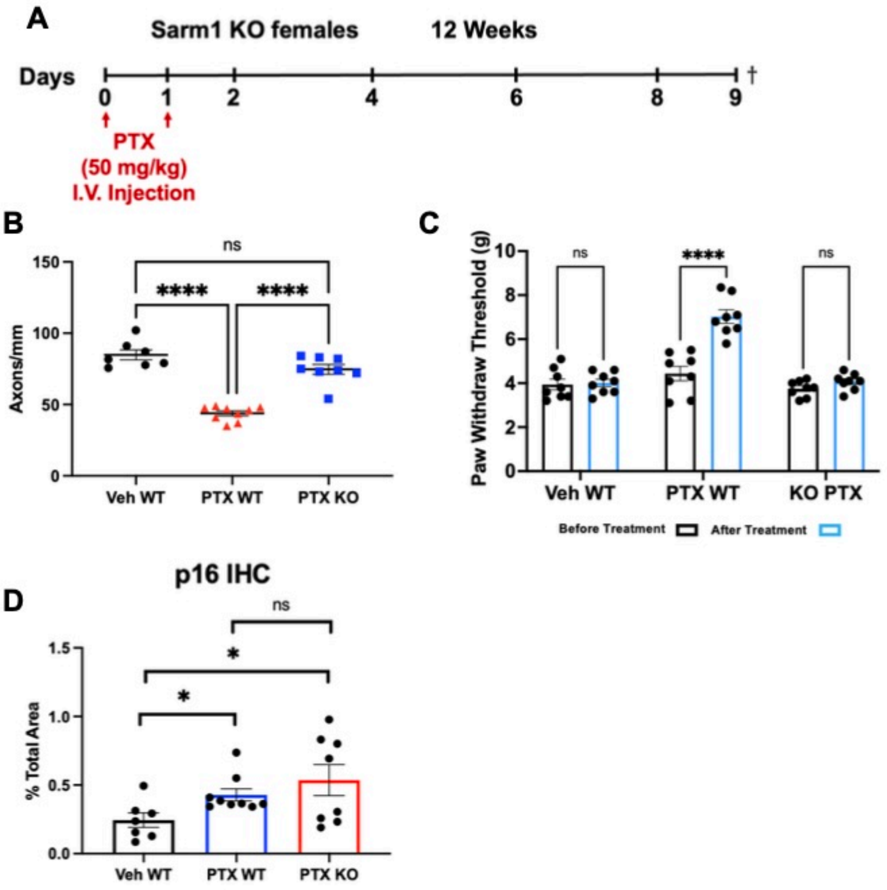


**Extended Data Fig. 4:**

### Ex.t Fig. 5

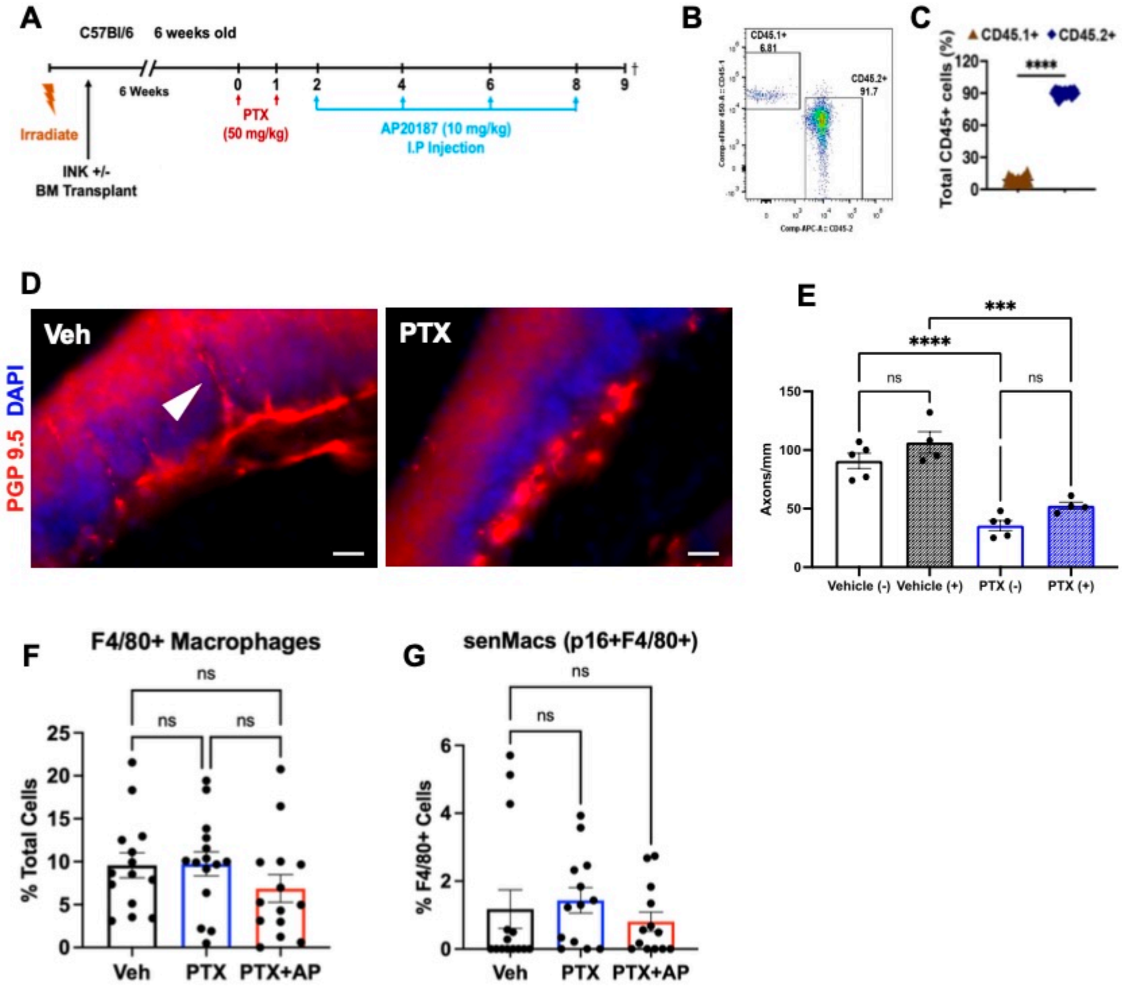


Extended Data Fig. 5:

### Ex.t Fig. 6

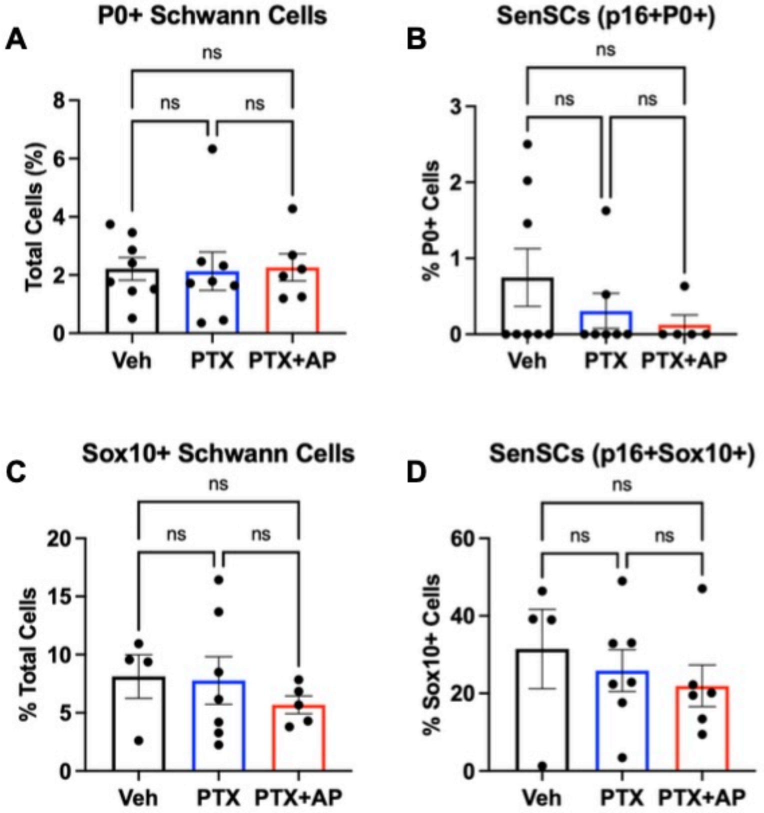


**Extended Data Fig. 6:**

### Ex.t Fig. 7

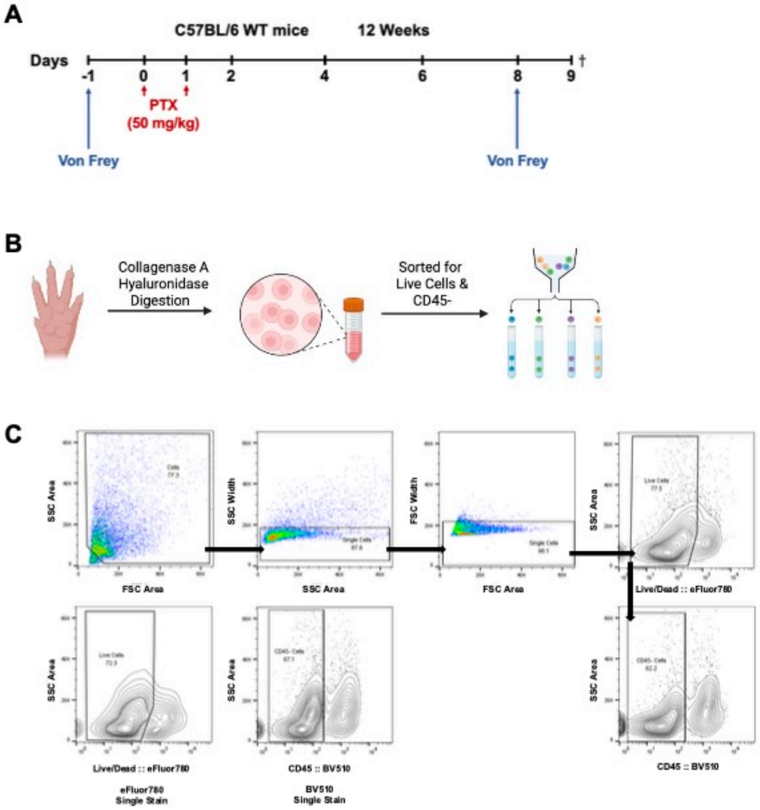


**Extended Data Fig. 7:**

### Ex.t Fig. 8

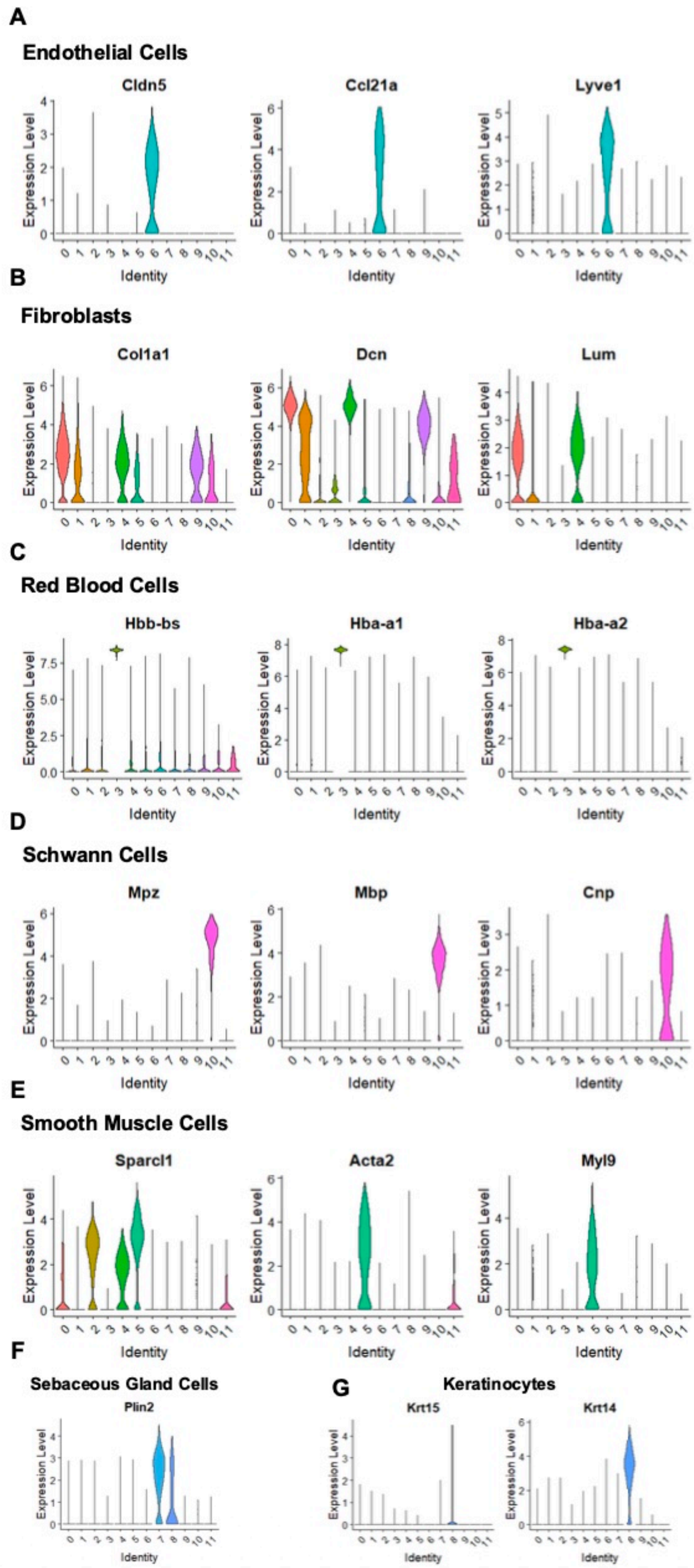


**Extended Data Fig. 8:**

### Ex.t Fig. 9

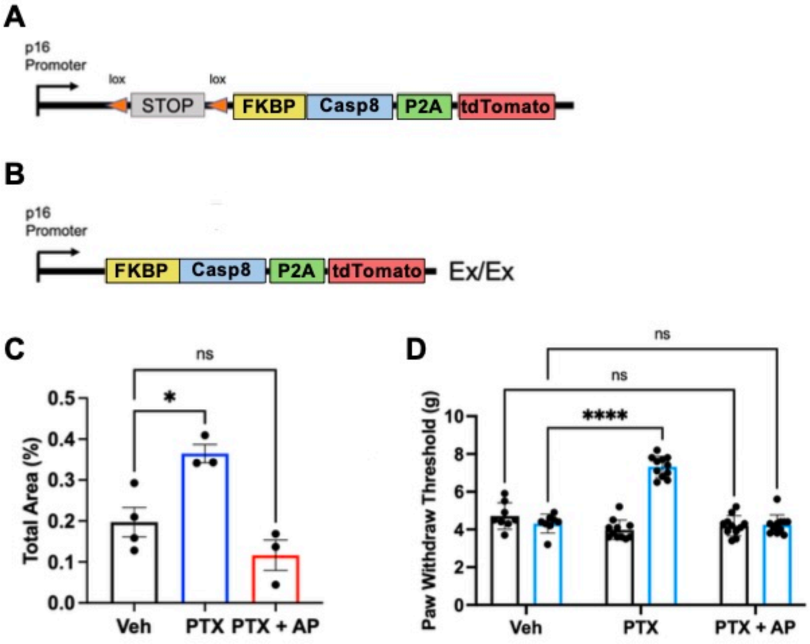


**Extended Data Fig. 9:**

### Ex.t Fig. 10

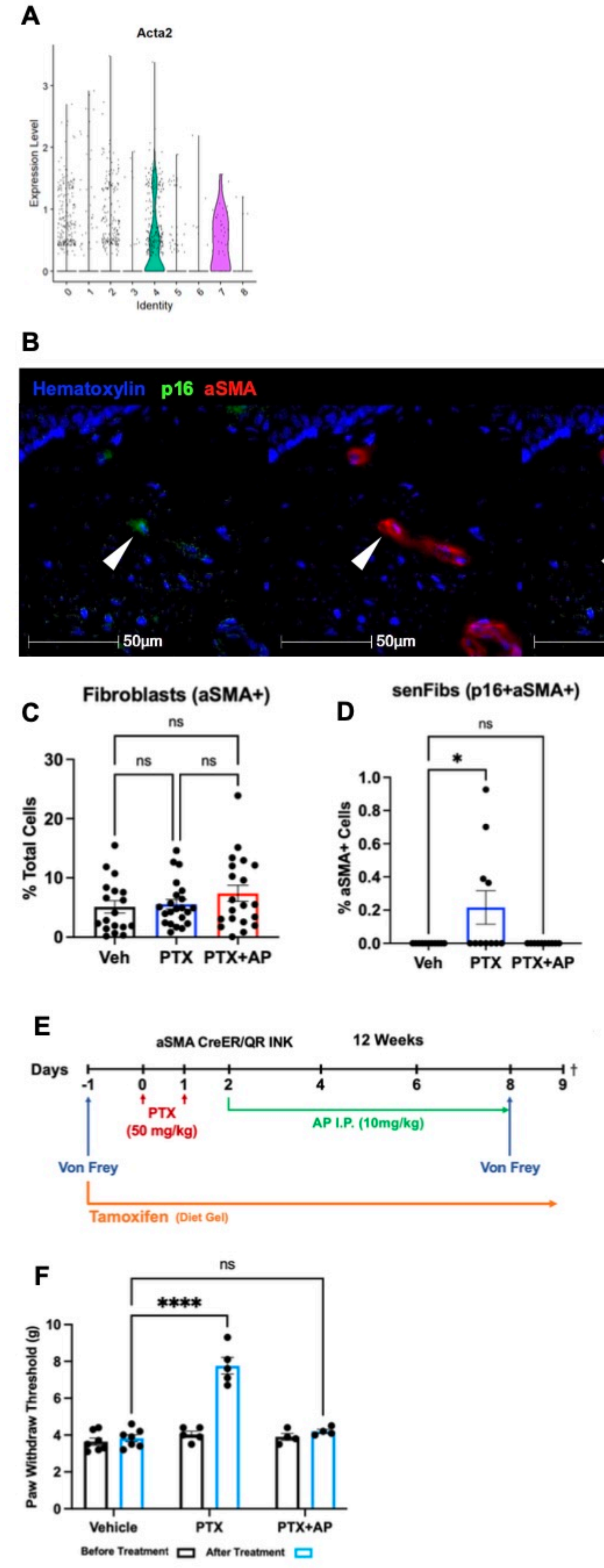


**Extended Data Fig. 10:**

### Ex.t Fig. 11

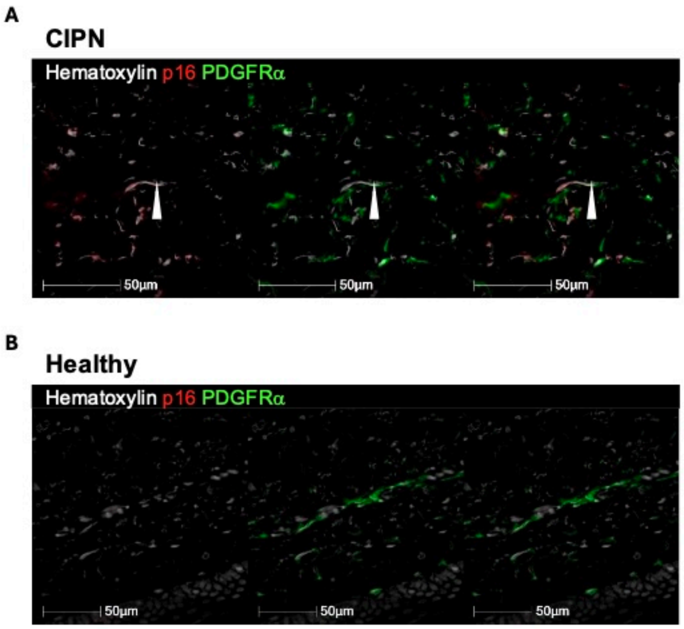


##### Extended Data Fig. 11:

### Ex.t Fig. 12

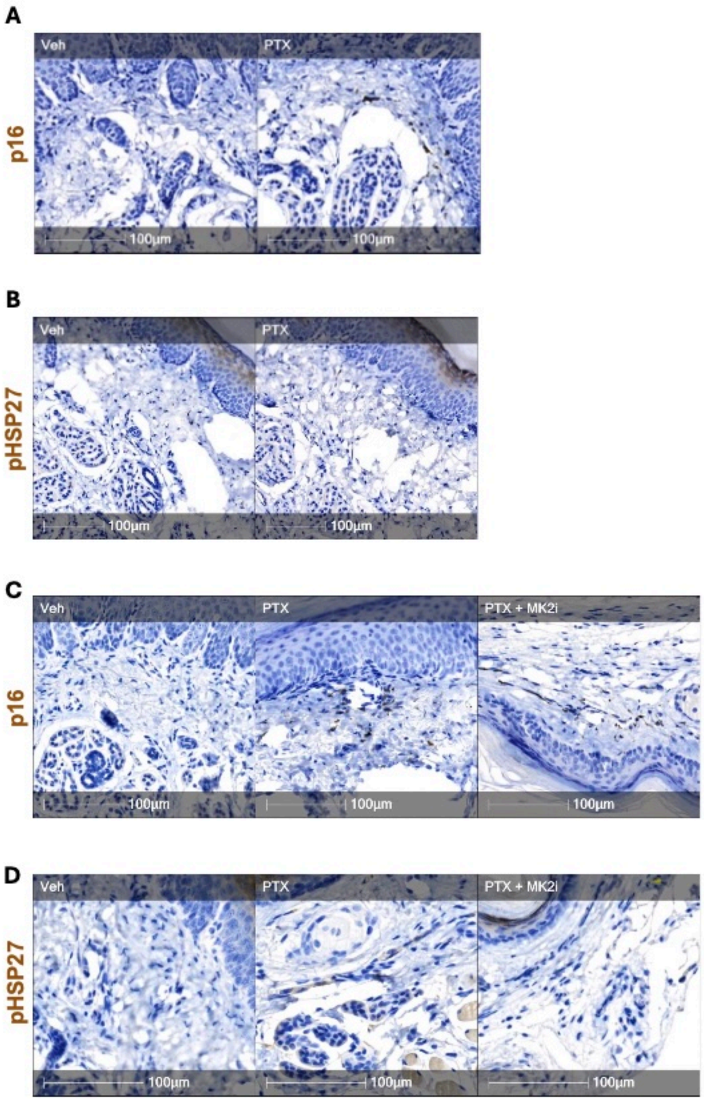


**Extended Data Fig. 12:**

### Ex.t Fig. 13

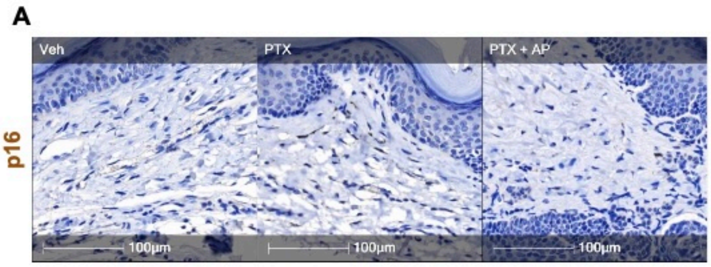


**Extended Data Fig. 13:**
