## Supplementary material for "Senescent peripheral fibroblasts initiate chemotherapy-induced peripheral neuropathy": Supple figures legends

**EXTENDED DATA FIGURES**

**Extended Data Fig. 1: Genetic elimination of senescent cells prevents CIPN in male INK mice.**

**(A**) Quantification of innervating axons from male INK mice; Veh (n=4), Veh+AP (n=4), PTX (n=4), and PTX+AP (n=3). One-way ANOVA was performed, data are represented as mean ± SEM. ****p< 0.001; ns, not significant.

**Extended Data Fig. 2: Senolytic elimination of senescent cells prevents CIPN in cisplatin treated mice.**

(A) Schema: DQ treatment schedule for 12-week-old WT mice treated with Veh or Cisplatin (2.3mg/kg) I.P. injection. **(B)** Quantification of innervating axons from cisplatin treatment; Veh (n=4), Cisplatin (n=5), Cisplatin+DQ (n=4). One-way ANOVA was performed, data are represented as mean ± SEM. ****p< 0.001; ns, not significant. **(C)** Von Frey results on Day -1 (black) and Day 8 (blue) from same mice shown in B; (2-way ANOVA was performed, data are represented as mean ± SEM. ****p <0.0001; ns, not significant). **(D)** Quantification of IHC for p16^+^ area out of total tissue area from same mice shown in B. One-way ANOVA was performed, data are represented as mean ± SEM. *p< 0.05; ns, not significant.

**Extended Data Fig. 3: DRGs, sural and sciatic nerves show no changes in senescence markers.**

**(A)** Representative IHC images for p16 (brown) in DRG sections from 12-week-old WT mice treated with Veh (n=3) or PTX (n=3). Red arrowheads indicate p16^+^ cells. **(B)** Quantification of p16^+^ area per total tissue area from same mice shown in A. Two-tail Student t-test was performed; data are represented as mean ± SEM. ns, not significant. **(C)** Representative IF images for SPiDER (green) and DAPI (blue) in DRG sections from 12-week-old WT mice treated with Veh (n=3) or PTX (n=3). **(D)** Quantification of SPiDER^+^ cells per total tissue area from same mice shown in C. Two-tail Student t-test was performed; data are represented as mean ± SEM. ns, not significant. **(E)** Representative IHC images for p16 (brown) in sciatic nerve sections from 12-week-old WT mice treated with Veh (n=5) or PTX (n=5). Red arrowheads indicate p16^+^ cells. **(F)** Quantification of p16^+^ area per total tissue area from same mice shown in E. Two-tail Student t-test was performed; data are represented as mean ± SEM. ns, not significant. **(G)** Representative IHC images for p16 (brown) in sural nerve sections from 12-week-old WT mice treated with Veh (n=5) or PTX (n=5). Red arrowheads indicate p16^+^ cells. **(H)** Quantification of p16^+^ area per total tissue area from same mice shown in G. Two-tail Student t-test was performed; data are represented as mean ± SEM. ns, not significant.

**Extended Data Fig. 4: PTX induces senescence in SARM1 KO animals.**

**(A) Schema:** PTX treatment delivery in 12-week-old SARM1 KO mice. **(B)** Quantification of innervating axons from SARM1 KO study; WT Veh (n=7), WT PTX (n=9), SARM1 KO PTX (n=8). One-way ANOVA was performed to compare innervating axon numbers, data are represented as mean ± SEM. ****p<0.0001. **(C)** Von Frey results on Day -1 (black) and Day 8 (blue) from same mice shown in B; (2-way ANOVA was performed to compare behavioral outputs, data are represented as mean ± SEM. ***p <0.001; ns, not significant). **(D)** Quantification of p16^+^ area per total tissue area from same mice shown in B. One-way ANOVA was performed; data are represented as mean ± SEM. ns, not significant.

**Extended Data Fig. 5: Bone marrow derived immune cells and tissue resident macrophages do not drive CIPN.**

**(A) Schema:** Bone marrow transplant timeline with PTX and AP treatment details. **(B)** Flow cytometry gating for CD45.1 and CD45.2 percentages. **(C)** Quantification of successful engraftment as previously shown in Raut et al. [31]. **(D)** Representative IF images for PGP9.5 (red) and DAPI (blue) in hind paw sections from 12-week-old BM transplant mice treated with Veh INK^-^ (n=5), Veh INK^+^ (n=5), PTX INK^-^ (n=5) or PTX INK^+^ (n=5). White arrowheads indicate innervating axon. **(E)** Quantification of innervating axons from same mice shown in D; (One-way ANOVA was performed, data are represented as mean ± SEM. ***p<0.001; ****p< 0.0001; ns, not significant). **(F)** mIHC quantification for F4/80^+^ macrophages out of total cells in INK mice. Veh (n=8), PTX (n=8), PTX+AP (n=6). One-way ANOVA was performed, data are represented as mean ± SEM. ns, not significant. **(G)** mIHC quantification for p16^+^F4/80^+^ senescent macrophages out of total macrophages in INK mice. Veh (n=8), PTX (n=8), PTX+AP (n=6). One-way ANOVA was performed, data are represented as mean ± SEM. ns, not significant.

**Extended Data Fig. 6: Senescent Schwann cells do not drive CIPN.**

**(A)** mIHC quantification for P0^+^ myelinating Schwann cells out of total cells in INK mice. Veh (n=8), PTX (n=8), PTX+AP (n=6). One-way ANOVA was performed, data are represented as mean ± SEM. ns, not significant. **(B)** mIHC quantification for p16^+^P0^+^ senescent myelinating Schwann cells out of total P0^+^ Schwann cells in INK mice. Veh (n=8), PTX (n=7), PTX+AP (n=6). One-way ANOVA was performed, data are represented as mean ± SEM. ns, not significant. **(C)** mIHC quantification for SOX10^+^ Schwann cells out of total cells in INK mice. Veh (n=4), PTX (n=7), PTX+AP (n=5). One-way ANOVA was performed, data are represented as mean ± SEM. ns, not significant. **(D)** mIHC quantification for p16^+^SOX10^+^ senescent Schwann cells out of total SOX10^+^ Schwann cells in INK mice. Veh (n=4), PTX (n=7), PTX+AP (n=6). One-way ANOVA was performed, data are represented as mean ± SEM. ns, not significant.

**Extended Data Fig.7: Single cell timeline, cell preparation and gating.**

**(A) Schema:** PTX treatment delivery in 12-week-old WT mice used for single cell RNA sequencing. **(B)** Schema: Tissue digestion and cell preparation for sorting. **(C)** Gating strategy for sorting cell populations.

**Extended Data Fig. 8: Cell markers for scRNA sequencing clustering.**

**(A)** Violin plots for endothelial cell markers including *Cldn5*, *Ccl21a*, and *Lyve1*. **(B)** Violin plots for fibroblast markers including *Col1a1*, *Dcn*, and *Lum*. **(C)** Violin plots for red blood cell markers including *Hbb-bs*, *Hba-a1*, and *Hba-a2*. **(D)** Violin plots for Schwann cell markers including *Mpz*, *Mbp*, and *Cnp*. **(E)** Violin plots for smooth muscle cell markers including *Sparcl1*, *Acta2*, and *Myl9*. **(F)** Violin plot for sebaceous gland cell markers including *Plin2*. **(G)** Violin plots for keratinocyte markers including *Krt15* and *Krt14*.

**Extended Data Fig. 9: QR INK-ATTAC Construct and QR Ex Validation.**

**(A)** QR INK-ATTAC construct schematic. **(B)** QR^+/+^ mouse construct schematic. **(C)** Quantification of p16^+^ area per total tissue area from QR^+/+^ mice Veh (n-4), PTX (n=3), PTX+AP (n=3). One-way ANOVA was performed; data are represented as mean ± SEM. *p>0.05; ns, not significant. **(D)** Von Frey results on Day -1 (black) and Day 8 (blue) from QR^+/+^ experiments; 2-way ANOVA was performed to compare behavioral outputs, data are represented as mean ± SEM. ****p <0.0001; ns, not significant.

**Extended Data Fig. 10: aSMA+ senescent fibroblasts drive CIPN.**

**(A**) Violin plot of αSMA expression across all fibroblast clusters. **(B)** mIHC were performed on hind paw samples of 12-week-old INK-ATTAC mice using p16 (green) and αSMA (red) specific antibodies and nuclei were counterstained with hematoxylin and shown as blue. Cells double positive for p16 and αSMA are denoted by white arrowheads. Individual stains were pseudo colored as indicated and then merged (last panel). **(C)** mIHC quantification for αSMA ^+^ fibroblasts out of total cells in INK mice. Veh (n=12), PTX (n=15), PTX+AP (n=12). One-way ANOVA was performed, data are represented as mean ± SEM. ns, not significant. **(D)** Quantification of senescent fibroblasts (p16+ αSMA^+^) out of total fibroblasts (αSMA^+^). One-way ANOVA was performed, data are represented as mean ± SEM. ns, not significant. **(E)** Schema: Tamoxifen and AP delivery for αSMA CreER QR mice that received either Veh or PTX. **(F)** Von Frey results on Day -1 (black) and Day 8 (blue) from same αSMA CreER QR mice; (2-way ANOVA was performed, data are represented as mean ± SEM. ****p <0.001; ns, not significant).

**Extended Data Fig. 11: Human biopsy mIHC representative images.**

**(A)** mIHC were performed on human skin punch biopsies from healthy patients or patients with CIPN (shown) using p16 (red) and PDGFRa (green) specific antibodies and nuclei were counterstained with hematoxylin and shown as white. Cells double positive for p16 and PDGFRα are denoted by white arrowheads. Individual stains were pseudo colored as indicated and then merged (last panel). **(B)** mIHC were performed on human skin punch biopsies from healthy patients (shown) or patients with CIPN using p16 (red) and PDGFRa (green) specific antibodies and nuclei were counterstained with hematoxylin and shown as white. Cells double positive for p16 and PDGFRα are denoted by white arrowheads. Individual stains were pseudo colored as indicated and then merged (last panel).

**Extended Data Fig. 12: Representative images for MK2 studies.**

**(A)** Representative IHC images for p16 (brown) in hind paw sections from 12-week-old MK2 KO mice treated with Veh or PTX. **(B)** Representative IHC images for pHSP27 (brown) in hind paw sections from 12-week-old MK2 KO mice treated with Veh or PTX. **(C)** Representative IHC images for p16 (brown) in hind paw sections from 12-week-old WT mice treated with Veh, PTX, or PTX+MK2i. CDD2231 (MK2i) was used for these studies. **(D)** Representative IHC images for pHSP27 (brown) in hind paw sections from 12-week-old WT mice treated with Veh, PTX, or PTX+MK2i. CDD2231 (MK2i) was used for these studies.

**Extended Data Fig. 13: Representative images for rescue studies.**

**(A)** Representative IHC images for p16 (brown) in hind paw sections from 12-week-old INK-ATTAC mice treated with Veh, PTX, or PTX+AP.
